# VIP36 preferentially binds to core-fucosylated N-glycans: a molecular docking study

**DOI:** 10.1101/092460

**Authors:** Klaus Fiedler

## Abstract

Alpha 1-6 fucosyltransferase (Fut8) is known for its properties as an enhancer of nonsmall cell lung cancer metastasis and as a suppressor in hepatocellular carcinoma cells (Hep3B). Promising candidates of affected molecules include E-cadherin. In its absence, during epithelial-mesenchymal transition, the pathway triggers signaling to the nucleus via β-catenin-TCF/LEF. Contrarily, in less metastatic tumors, Fut8 stimulates cell-cell adhesion. Regulated classes of molecules could also include the sorting machinery of polarized epithelial cells, sorted ligands or both, that may be altered in cellular transformation. I have analyzed here the cargo receptor VIP36 (Vesicular-integral membrane protein of 36 kD) for carbohydrate interaction. It has been described as a lectin in the ERGIC (endoplasmic reticulum-Golgi intermediate compartment), Golgi apparatus and plasma membrane. The docking reveals top-interacting carbohydrates of the N-glycan and O-glycan class that encompass N-linked glycans of high mannose and equally complex type which likely function as sorted ligands in epithelial cells. O-glycans score lower and include core 2 residue binding. I show that fucose core modifications by Fut8 stimulate binding of N-linked glycans to VIP36, which is known to be different from binding of galectins 3 and 9. This suggests that Fut8-upregulation may directly alter the affinity of sorted cargo and may enhance the sorting to the apical pathway as exemplified in hepatocytes and traffic to bile. High affinity binding of the ganglioside GM1 carbohydrate headgroup to VIP36 suggests a linkage with protein and glycosphingolipid apical transfer in epithelial cells. Thus, this fundamental approach with large scale docking of 165 carbohydrates including 19 N-glycan high mannose, 17 Nglycan hybrid, 9 N-glycan complex, 17 O-glycan core, 27 Sialoside, 25 Fucoside and 51 other glycan residues suggests, that linked cargo-receptor apical transport may provide a path to epithelial polarization that may be modulated by core fucosylation.

## Background

VIP36 was identified as a component of post-Golgi secretory vesicles (Fiedler et al 1994) and had been shown to be involved in sorting of glycosylated alpha-amylase in the exocytic pathway (Hara-Kuge et al 2002; Hara-Kuge et al 2003; Arshad et al 2013). In polarized epithelia (Mellman and Nelson 2008; Rodriguez-Boulan and Macara 2014), it had been established, that glycosylation of growth hormone was important for apical sorting (Scheiffele et al 1995); erythropoietin and apical membrane proteins were also found to require glycosylation for apical targeting (Kitagawa et al 1994; Gut et al 1998). The glycan affinities to VIP36 had been a matter of discussion.

The VIP36 protein is similar to ERGIC-53 (Itin et al 1995) and belongs to the class of eukaryotic leguminous-type lectins. It is abundantly expressed in mouse kidney, liver, intestine, lung and spleen (Fiedler and Simons 1996). In humans, it is abundantly present in testis, epididymis, seminal vesicles, placenta and thyroid gland, and also expressed in liver, gallbladder, pancreas, stomach, cervix, endometrium, fallopian tube, bone marrow, lymph node, tonsil, brain and lung (Uhlén et al 2015). In computational analyses it had been demonstrated to be related to the plant lectin in *Bauhinia purpurea* (Fiedler et al 1994; Fiedler and Simons 1994). Another lectin, galectin-9, has recently been implicated in apical sorting and was found to influence the cell polarity of MDCK (Madin-Darby Canine Kidney) cells (Mishra et al 2010). The relationship of both lectins, if any, remains an open question. Galectins 3 and 4 have also previously been shown to be involved in apical transport in MDCK and HT-29 (enterocyte-like) cells, respectively (Delacour et al 2006). In particular galectin-3 which is localized to the cell surface, the Golgi and the cytoplasm is found to play a role in tumor metastasis (Takenaka et al 2002). This protein is found as a binding partner of β-catenin-TCF/LEF which co-localizes to the nucleus of cells and stimulates transcriptional activity (Shimura et al 2004). Is it plausible that several lectins are involved in generating the epithelial lipid and protein polarity that leads to physiological vectorial transport while alternatively stimulating cancer progression? Wnt secretion, the molecular ligand of the Wnt/frizzled-pathway, involves N-linked glycosylation and has recently been shown to benefit from complex glycosylation (Wnt11) in the apical transport, and surprisingly may require galectins and Opossum, a p24 protein, for secretion (Fiedler et al 1996; Buechling et al 2011; Yamamoto et al 2013).

The glycoproteostasis involves ER enzymes and lectins calreticulin (CRT) and calnexin (CNX) that serve to assist the ordered glycoprotein folding and quality control and ER-associated protein degradation (ERAD)(Helenius and Aebi 2001; Satoh et al 2010; Hebert et al 2014). VIP36 had previously been demonstrated to be localized to the Golgi apparatus, exocytic carrier vesicles and the cell surface and lends itself to a sorting function in the Golgi complex (Fiedler et al 1994). The endogenous protein was then also visualized in punctate structures that had been co-localized with the ER-Golgi intermediate compartment (ERGIC) constituent ERGIC-53. The labelling of COPI or sialyl-transferase of the trans-Golgi cisterna did further demonstrate substantial overlap as shown in MDCK cells (Fullekrug et al 1999). In contrast to MDCK cells, where in EM the Golgi apparatus and the plasma membrane were preferentially decorated with anti-VIP36 antibodies (Hara-Kuge et al 2002) it was found in other epithelial cells, that, moreover, endocytic structures and late endosomes were labeled (K. Fiedler and K. Simons, unpublished results). Immature granules and mature secretory granules as well as the trans-Golgi apparatus were decorated with antibodies in parotid secretory cells as seen in EM (Shimada et al 2003).

VIP36 and ERGIC-53 contain a single carbohydrate recognition domain (CRD). The D1 arm of mannoses linked to Asn-N-sequons was found to form extensive hydrogen bonds with VIP36 in the crystal 2DUR (Satoh et al 2007) and interestingly, the coordination of Ca^2+^ was found to orient the side chains that had previously been speculated to be involved in carbohydrate and calcium binding based on this similarity (Fiedler et al 1994), i.e. Asp131 and Asn166 as well as His190. Some variation may be introduced by the largest variability of protein sequences of VIP36s and related proteins in the glycan and calcium-binding loop that we described as region II (Fiedler et al 1994). Next to conserved amino acids (Asn166 in VIP36 and ERGIC-53, VIPL/LMAN2-L; not in mouse or human ERGL/LMAN1-L and yeast EMP46-like; Table S3)(see (Neve et al 2003)) in representative members this shows gaps. Although the loops are usually not ascribed particular functions, it could be that the large evolutionary glycan variation may be adapted by amino acid residues in the vicinity of the binding site. Galectin-9 contains two CRDs and the N-terminal CRD was described to interact with polylactosamines and some glycosphingolipid (GSL) headgroups (Hirabayashi et al 2002; Nagae et al 2008; Nagae et al 2009; Mishra et al 2010). An activity as an eosinophil chemoattractant had also been discovered (Sato et al 2002). A link of cargo receptors to specific bilayer lipids was also described: it was found that sphingomyelin 18 interacted with the transmembrane domain of the p24 protein and regulated ER to Golgi transport (Fiedler and Rothman 1997; Contreras et al 2012). The critical glutamine residue contained within the bound signature sequence was previously found to halt export of p24 chimeras in the early secretory pathway once mutated to alanine.

Here the large scale screening of carbohydrates by computational docking (Trott and Olson 2010) replaces the glycan screens usually developed for microarrays or mass spectrometry which due to technical difficulties do usually not generate nearly complete solutions. The present data may contribute to shed light on the role of fucose in carbohydrate N-glycan binding of the L-type lectins and possible indications of branch preference of GlcNAc-transferases with respect to apical sorting of glycoproteins. The results from cognate docking suggest that GlcNAc-Transferase V (GlcNAcT-V) modification of the α6 mannose arm of N-glycan chains is not of high affinity, however, the GlcNAcT-IV modified glycans with GlcNAc added to the α3 mannose arm may represent the preferred high affinity cargo.

N-glycan chains are modified by adding fucose residues to their terminal chains (fucosyltransferase 2; Fut2) or to the N-glycan core (fucosyltransferase 8; Fut8). Recent discovery of the influence of Fut2 activity on the protective role of fucosylation suggests a function of terminal modifications in host-microbial interactions (Pickard et al 2014; Goto et al 2014) whereas core fucosylation by Fut8 has been described as possible sorting signal in protein traffic in hepatocytes to the bile duct (Nakagawa et al 2006). Fut8 was, furthermore, shown to regulate metastasis in nonsmall cell lung cancer (NSCLC) but overexpression in hepatoma cells (Hep3B) was demonstrated to suppress intrahepatic metastasis in athymic mice (Miyoshi et al 1999; Chen et al 2013). The E-cadherin-β-catenin-TCF/LEF pathway and α5β1 integrin have been invoked in these analyses. The relation and intricate balance of cell-cell and cell-extracellular matrix adhesion to protein fucosylation (and other glycan modifications) may explain the latter two findings and glycan modifications with respect to protein traffic and polarity of epithelia.

## Results

### Lectins

Next to legume L-type lectins, CNX, CRT, Emp47, pentraxins (C-reactive protein, serum amyloid P) and vp4 of rotavirus also belong to L-type lectins (Fiedler and Simons 1994; Neve et al 2003; Etzler et al 2009). An overview of a structural comparison is shown in Table 1. Galectins, although only up to 10% identical in amino acid sequence, show a similar fold to VIP36. The carbohydrate binding site is, yet, removed from the site of ERGIC-53 and VIP36 glycan interaction. VIP36 an ERGIC-53 are 30.4% identical in sequence within the L-type lectin domain. Galectin-3 is only 10.1% identical in sequence to VIP36, and galectins do not contain a calcium binding site and coordinate glycans on the large β-sheet, i.e. not via loops (see Fig. 3, 7). In the present analysis I have restricted the molecular docking to VIP36.

**Table I.**
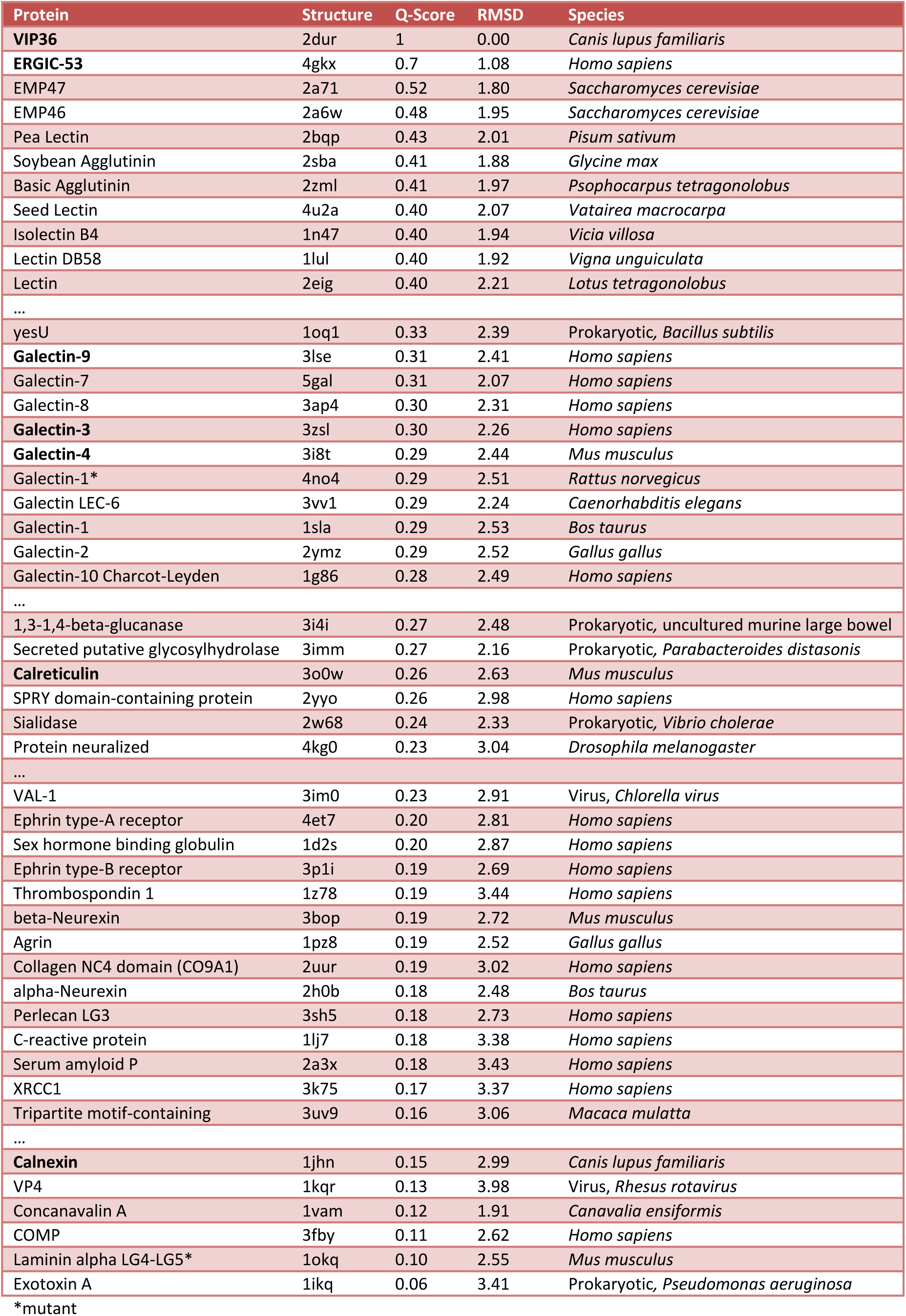
Structural comparison of leguminous-type lectins. VIP36 (2DUR) was used in the database search with PDB. Structural hits were ranked according to PDBeFold **Q-Score**. **RMSDs**, each **Structure** found and the **Protein** name is indicated. All retrieved structures were derived from eukaryotic proteins unless otherwise indicated (**Species**), some proteins were crystallized in mutant form and top-scoring protein isoforms of each entry were given. The results were collated from 2860 structure files; the lowest match was adjusted to 60%. Entries that had previously been ranked and determined to be structurally similar to L-type lectins (1JHN, 3FBY, 1IKQ) were added for non-inclusive ranking. Proteins shown in bold are discussed in the text.

PDBeFold search with 2DUR
| Protein | Structure | Q-Score | RMSD | Species |
| --- | --- | --- | --- | --- |
| **VIP36** | 2dur | 1 | 0.00 | *Canis lupus familiaris* |
| **ERGIC-53** | 4gkx | 0.7 | 1.08 | *Homo sapiens* |
| EMP47 | 2a71 | 0.52 | 1.80 | *Saccharomyces cerevisiae* |
| EMP46 | 2a6w | 0.48 | 1.95 | *Saccharomyces cerevisiae* |
| Pea Lectin | 2bqp | 0.43 | 2.01 | *Pisum sativum* |
| Soybean Agglutinin | 2sba | 0.41 | 1.88 | *Glycine max* |
| Basic Agglutinin | 2zml | 0.41 | 1.97 | *Psophocarpus tetragonolobus* |
| Seed Lectin | 4u2a | 0.40 | 2.07 | *Vatairea macrocarpa* |
| Isolectin B4 | 1n47 | 0.40 | 1.94 | *Vicia villosa* |
| Lectin DB58 | 1lul | 0.40 | 1.92 | *Vigna unguiculata* |
| Lectin | 2eig | 0.40 | 2.21 | *Lotus tetragonolobus* |
| ... |  |  |  |  |
| yesU | 1oq1 | 0.33 | 2.39 | Prokaryotic, *Bacillus subtilis* |
| **Galectin-9** | 3lse | 0.31 | 2.41 | *Homo sapiens* |
| Galectin-7 | 5gal | 0.31 | 2.07 | *Homo sapiens* |
| Galectin-8 | 3ap4 | 0.30 | 2.31 | *Homo sapiens* |
| **Galectin-3** | 3zsl | 0.30 | 2.26 | *Homo sapiens* |
| **Galectin-4** | 3i8t | 0.29 | 2.44 | *Mus musculus* |
| Galectin-1* | 4no4 | 0.29 | 2.51 | *Rattus norvegicus* |
| Galectin LEC-6 | 3vv1 | 0.29 | 2.24 | *Caenorhabditis elegans* |
| Galectin-1 | 1sla | 0.29 | 2.53 | *Bos taurus* |
| Galectin-2 | 2ymz | 0.29 | 2.52 | *Gallus gallus* |
| Galectin-10 Charcot-Leyden | 1g86 | 0.28 | 2.49 | *Homo sapiens* |
| ... |  |  |  |  |
| 1,3-1,4-beta-glucanase | 3i4i | 0.27 | 2.48 | Prokaryotic, uncultured murine large bowel |
| Secreted putative glycosylhydrolase | 3imm | 0.27 | 2.16 | Prokaryotic, *Parabacteroides distasonis* |
| **Calreticulin** | 3o0w | 0.26 | 2.63 | *Mus musculus* |
| SPRY domain-containing protein | 2yyo | 0.26 | 2.98 | *Homo sapiens* |
| Sialidase | 2w68 | 0.24 | 2.33 | Prokaryotic, *Vibrio cholerae* |
| Protein neuralized | 4kg0 | 0.23 | 3.04 | *Drosophila melanogaster* |
| ... |  |  |  |  |
| VAL-1 | 3im0 | 0.23 | 2.91 | Virus, *Chlorella virus* |
| Ephrin type-A receptor | 4et7 | 0.20 | 2.81 | *Homo sapiens* |
| Sex hormone binding globulin | 1d2s | 0.20 | 2.87 | *Homo sapiens* |
| Ephrin type-B receptor | 3p1i | 0.19 | 2.69 | *Homo sapiens* |
| Thrombospondin 1 | 1z78 | 0.19 | 3.44 | *Homo sapiens* |
| beta-Neurexin | 3bop | 0.19 | 2.72 | *Mus musculus* |
| Agrin | 1pz8 | 0.19 | 2.52 | *Gallus gallus* |
| Collagen NC4 domain (CO9A1) | 2uur | 0.19 | 3.02 | *Homo sapiens* |
| alpha-Neurexin | 2h0b | 0.18 | 2.48 | *Bos taurus* |
| Perlecan LG3 | 3sh5 | 0.18 | 2.73 | *Homo sapiens* |
| C-reactive protein | 1lj7 | 0.18 | 3.38 | *Homo sapiens* |
| Serum amyloid P | 2a3x | 0.18 | 3.43 | *Homo sapiens* |
| XRCC1 | 3k75 | 0.17 | 3.37 | *Homo sapiens* |
| Tripartite motif-containing | 3uv9 | 0.16 | 3.06 | *Macaca mulatta* |
| ... |  |  |  |  |
| **Calnexin** | 1jhn | 0.15 | 2.99 | *Canis lupus familiaris* |
| VP4 | 1kqr | 0.13 | 3.98 | Virus, *Rhesus rotavirus* |
| Concanavalin A | 1vam | 0.12 | 1.91 | *Canavalia ensiformis* |
| COMP | 3fby | 0.11 | 2.62 | *Homo sapiens* |
| Laminin alpha LG4-LG5* | 1okq | 0.10 | 2.55 | *Mus musculus* |
| Exotoxin A | 1ikq | 0.06 | 3.41 | Prokaryotic, *Pseudomonas aeruginosa* |

\*mutant

### Cognate binding to VIP36

The intactness of carbohydrate structures was checked and confirmed. Structural docking was simulated with PyRx (Wolf 2009), utilizing the VIP36 2E6V and 2DUR structural data (Satoh et al 2007). 2E6V was initially analyzed in Global docking Fig. 1). Library residues of high mannose type bound with maximal affinities of up to −6.7 kcal/mol to docking sites, that in this global approach did not correspond to cognate docking. Only 13.6 % of all high mannose ligands showed at least one cognate docking site, and 4.4 % of high mannose ligands preferentially interacted with the authentic, apo-ligand binding pocket. None of the complex hybrid and only sialylated or fucosylated N-glycan termini, possibly of N-glycan complex type, bound preferentially in global docking, just as only some O-glycan cores would preferentially interact with the cognate binding pocket with considerate affinity. The global approach was thus abandoned and structures were, furthermore, used as energy minimized conformers generated by conjugate gradient minimization steps. The local approach resulted in 49%, 42% and 42% of all glycans interacting with the main binding pocket of 2DUR-A, 2DUR-B and 2E6V, respectively, at or better than −6.5 kcal/mol. In these models, a single calcium was preserved in the binding assay instead of adding a further calcium which would have only loosely bound by 3 coordinations in a similar position to the 2^nd^ calcium of the VIP36-like ERGIC-53 (Velloso et al 2003). The previously determined K_a_ of VIP36 for calcium in the exoplasmic domain was found to be approx. 2.5 × 10^4^ M^−1^ (Fiedler and Simons 1996) which is actually quite similar to the determined carbohydrate affinities in the range of 1 −2 × 10^4^ M^−1^. Ligands interacting at or better than −6.5 kcal/mol represented possible candidates for true binding site interaction.

**Figure 1.**
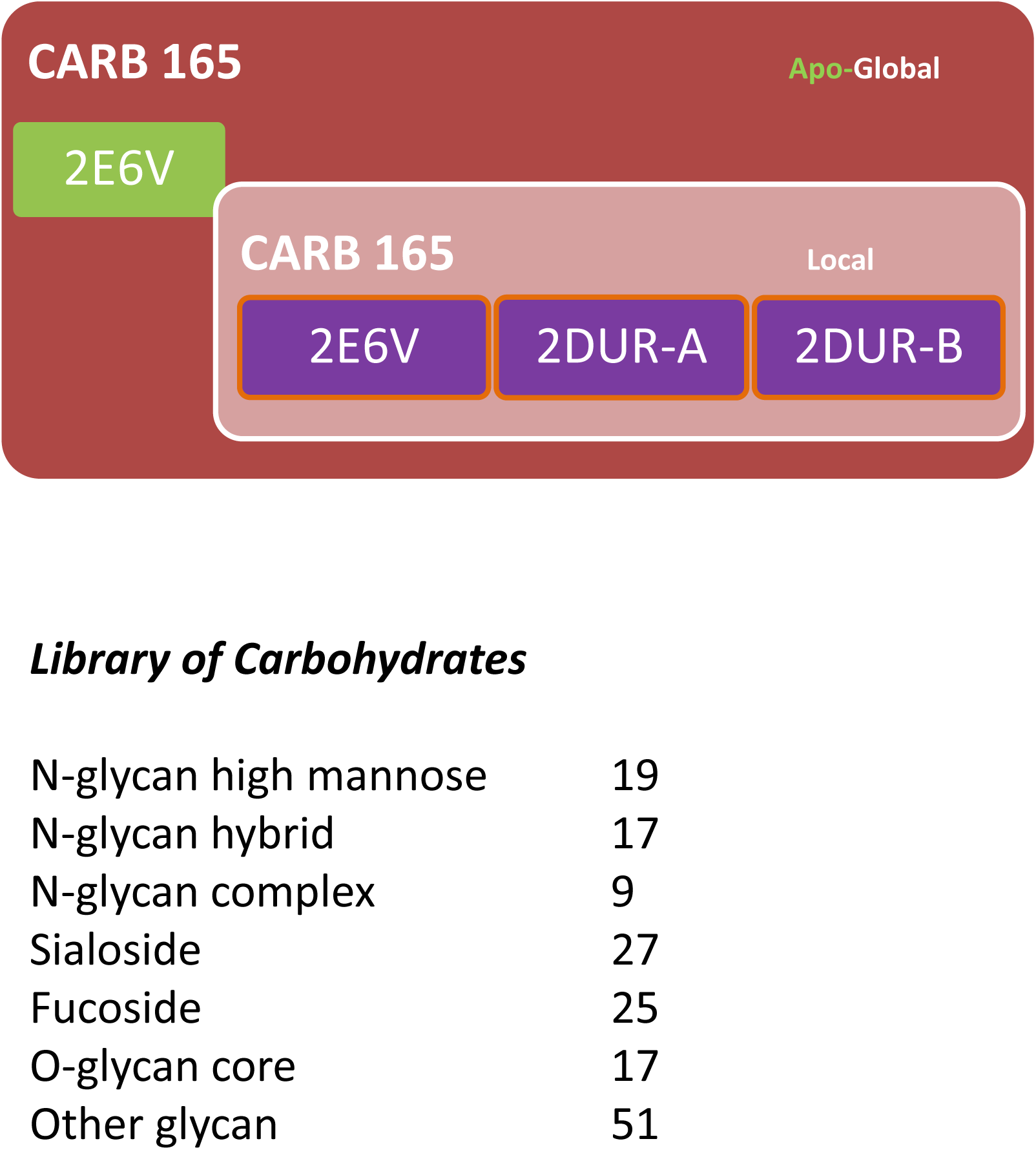
The CARB 165 library was constructed for this experimental approach. CARB 165 was tested against 2E6V, the structure of VIP36 chosen for a virtual ligand screen. 2E6V is established as “Apo” structure in “Global” screening, the structural studies resolved mannose residues in crystallization. The CARB 165 library was subsequently screened in the Local approach with 2DUR-A, 2DUR-B and towards binding to 2E6V, all three molecularly relaxed. “Holo enzyme” behavior is inferred from side chain rotation. The number of database entries in the respective carbohydrate group is indicated.

### Top scoring high-affinity interactors of VIP36 belong to the N-glycan, Ganglioside and O-glycan class

In order to fully explore the library Carb165 the 2DUR-A conformer was used. ViewDock was utilized to analyze H-bonds in between the modeled ligand and the holo-enzyme. The top scores of 15 Carb165 (Fig. S1, I-XV) glycans were presented according to motifs included Fig. 2); lacto-motifs were prevalent with five hits among the top 15. The interaction energies of raw, automated docking are graphed in Fig. S1 with top model numbers indicated that fulfill the structure test. The energies of these models differed slightly.

**Figure 2.**
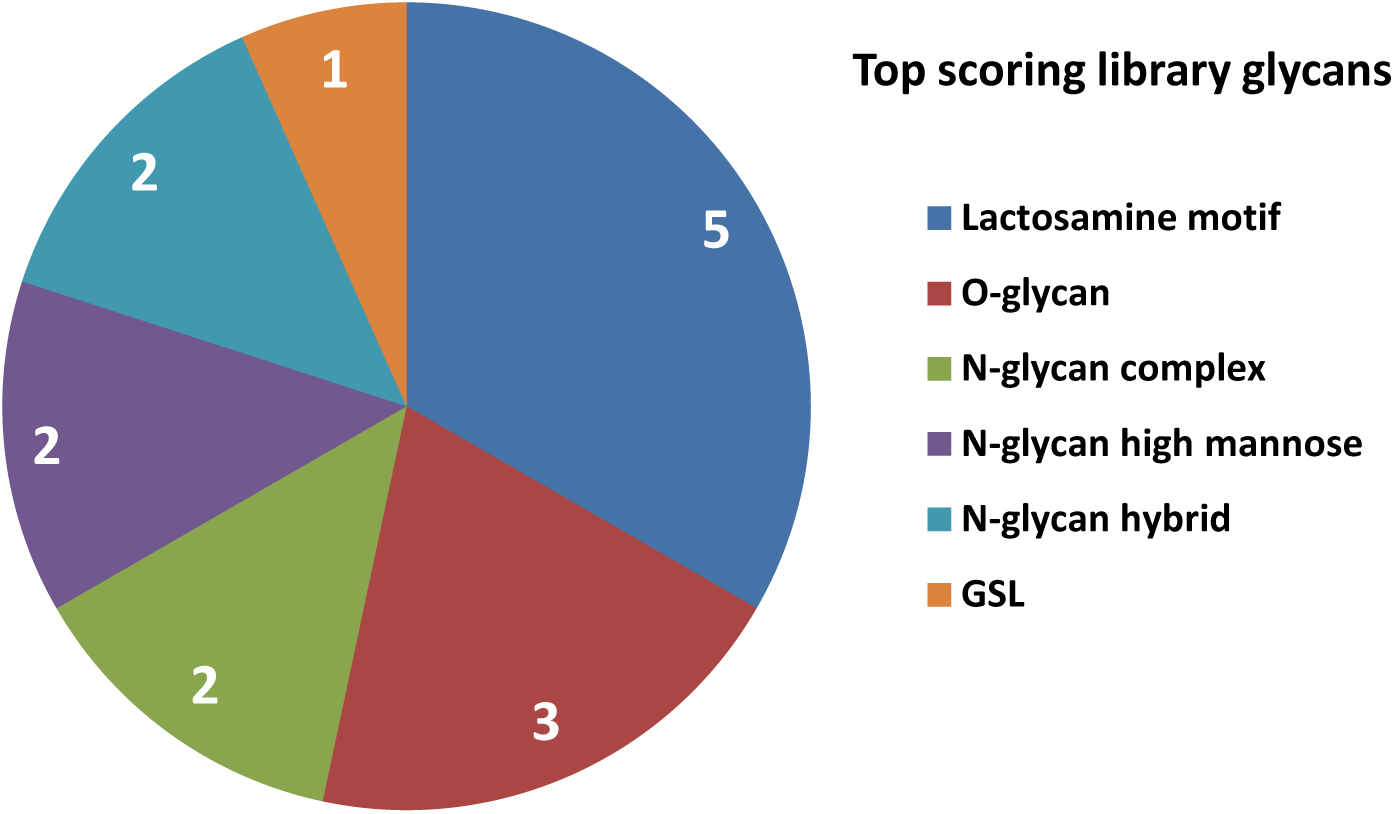
Summary of newly discovered glycan ligands of VIP36. The Carb165 library was screened for interactions with 2DUR-A. Number of glycan motifs among top scores are indicated (no double counts of extended structures or multiple motifs)(15 top scoring glycans).

The series of carbohydrates that bound with considerable, high affinity to Carb165 consisted of N-glycan high mannose and N-glycan core basic structures classified as complex glycans, followed by GM1 (Simons and Ikonen 1997; Sabourdy et al 2008), N-glycan high mannose and the lactosamine motif in the third rank. Using the structural criteria, the basic order of affinities does not change with the 3^rd^ ligand scoring number one amongst the top 15 (N-glycan high mannose) and the 13^th^ glycan scoring number two of the top selection (N-glycan core basic: complex and fucosylated). In the 9^th^ position of eligible carbohydrates extended O-glycan core 2 was found in rank number three using the structure tests (see Fig. S2; IX). Only the 9 top scoring models of each carbohydrate were tested. In a separate simulation by applying identical experimental parameters, β-D-GlcpNAc-(1-6)-[β-D-Galp-(1-3)]-α-D-GalpNAc-OH, the core 2 structure, as well as the T antigen, β-D-Galp-(1-3)-α-D-GalpNAc-OH, or Sialyl-Tn antigen, α-D-Neu5Ac-(2-6)-α-D-GalpNAc-OH was scoring less than the extended core, suggesting that the extensions of these O-glycans included relevant binding interactions.

The ganglio series GM1 (IV) was one of few GSL motifs in the entire database Carb165 (depleted in GSL residues). The GM1 headgroup is present, among other species, in *Campylobacter jejuni* as part of an epitope widely studied in human disease of autoimmune disorders caused by these bacteria (see Table S1). Further screening may later lead to precise determination of all glycosphingolipid affinities and binding specificities including their frequent H-bond acceptors and donors that may vary from other carbohydrate residues.

### Hydrogen-bonding of VIP36 to carbohydrate residues

The virtual screening approach was further used to determine the exact position of amino acids in VIP36 that would provide the bulk of hydrogen bond donor and acceptor activity. Docking in this gradient approach was described to function akin to “machine learned” binding (Trott and Olson 2010) applied to statistical physics Fig. 4). Results are summarized for measuring H-bonds in ViewDock with all 15 top scores of Carb165 against 2DUR-A. The overwhelming proximity of oxygens (3 Å zone distance) shows the large H-bond donor capacity of carbohydrate ligands with few nitrogens in molecular proximity of the VIP36 surface Fig. 3). The exact method used applies tolerances to H-bonding geometry that were found in crystallized structures (Mills and Dean 1996). The most probable ligand of carbohydrates in the VIP36 binding site is represented by the ND2 atom of Asn166. This ligand site is found within the central binding area at 11 o’clock and had previously been identified due to lectin homology of VIP36 to leguminous lectins (Lec_Baupu) (Fiedler et al 1994). This metal binding site appears to be conserved in leguminous lectins, ERGIC-53 and close VIP36 family members (L-type lectins). Above 41% of all 15.9 glycan model structures Fig. 3) generated by AutoDock form hydrogen bonds wherein the aforementioned residue act as donor. This was followed by Asp261 N and the Ser96 OG located both on the perimeter of the pocket in diametrically opposite position at the bottom of the central binding cavity. 14.4% and 10.0% of all carbohydrates use these as donors. The acceptor binding sites are more heterogeneous with five residues covering the largest share of 55.3% hydrogen-bond acceptors in total, the Pro146 O residue (13.1%) as seen on the upper right of the perimeter of the central indentation, the Asn166 O (11.6%) which is bidentate (Asn) located on the upper left side above the central trough, the Glu98 OE2 (10.9%) on the right of the binding pocket, the Thr259 O (10.2%) within the central indentation and Asp131 OD1 (9.5%) located just next to Asn166 on the upper perimeter of the central indentation (Fig. 4A). Large carbohydrates that bind to the central cavity form interactions using H-bond donors that cover an area of 700 Å^2^ coinciding with a modeled N-glycan of high mannose type (Satoh et al 2007). This was inserted in a nonflexible fashion and corresponds herein to topology 3 (Figs. 4B, S2).

**Figure 3.**
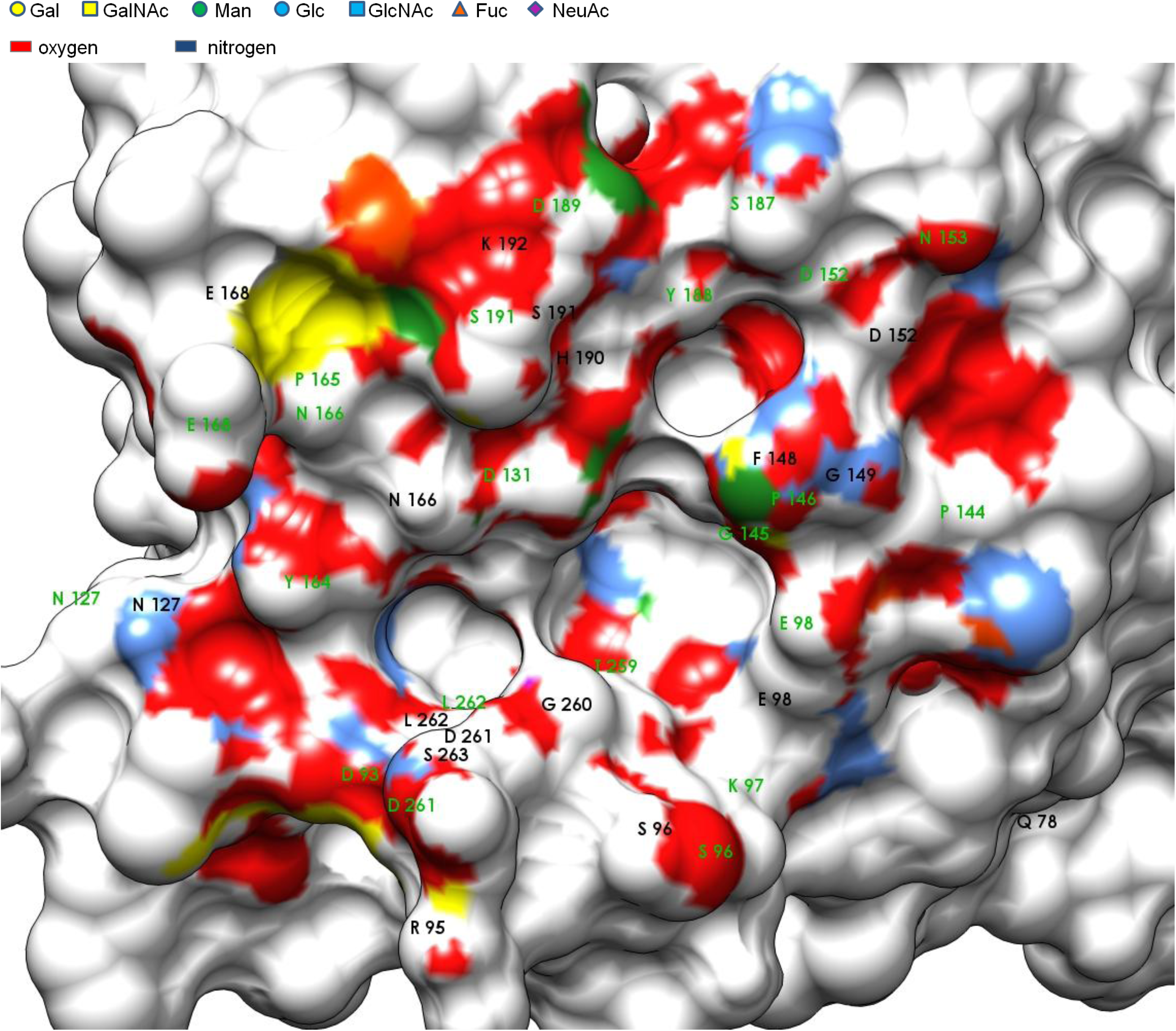
Carbohydrate interaction. Amino acids of VIP36 implicated in H-bonding to carbohydrate ligands are presented with surface-proximate groups, the legend indicates the IUPAC coloring of carbohydrates and their respective surface imprint at 3 Å. Few residues are covered behind E168. The binding site is shown with determined H-bond donors in black and acceptors in green for Carb165 (15.9 Models).

**Figure 4.**
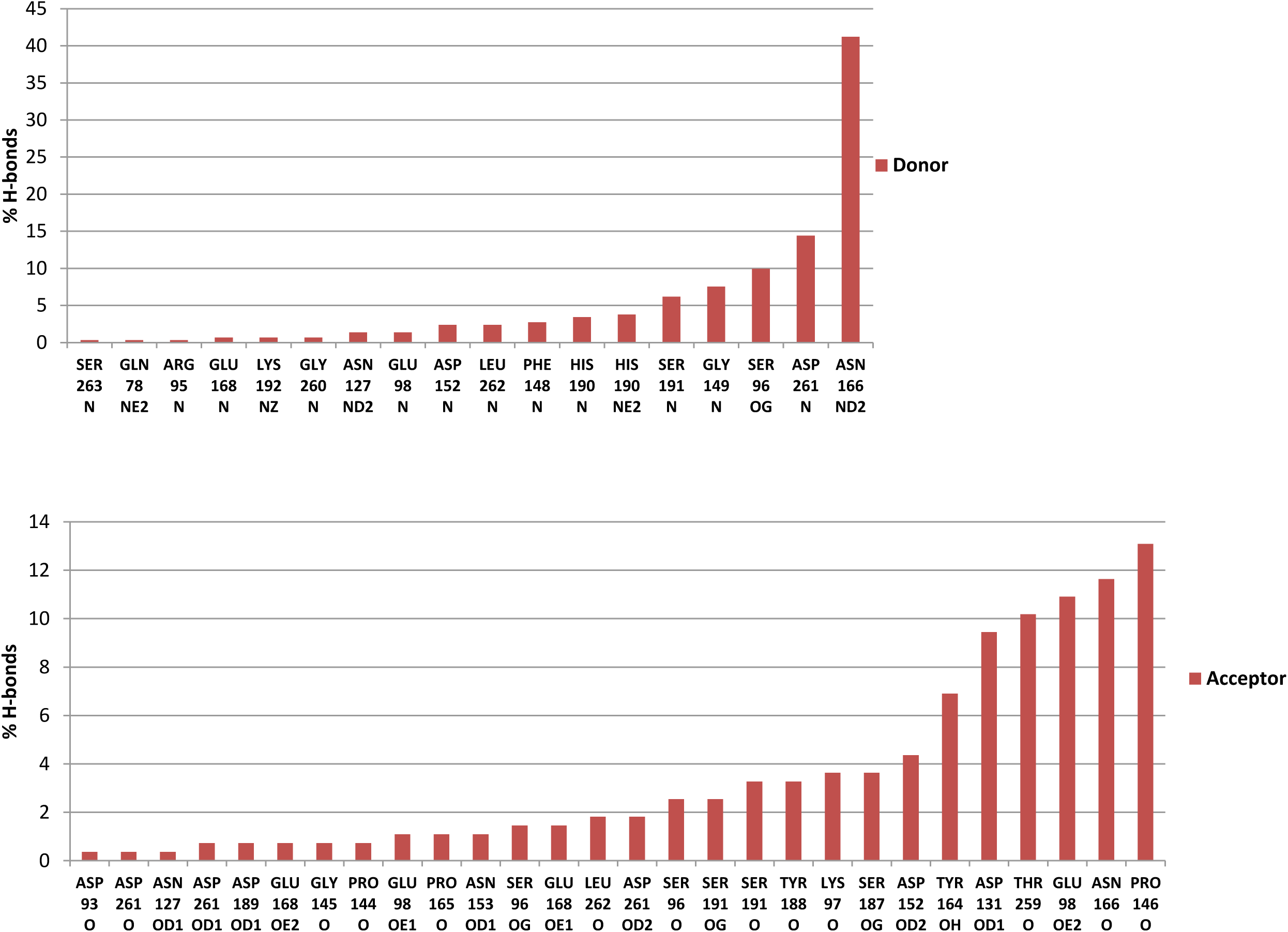
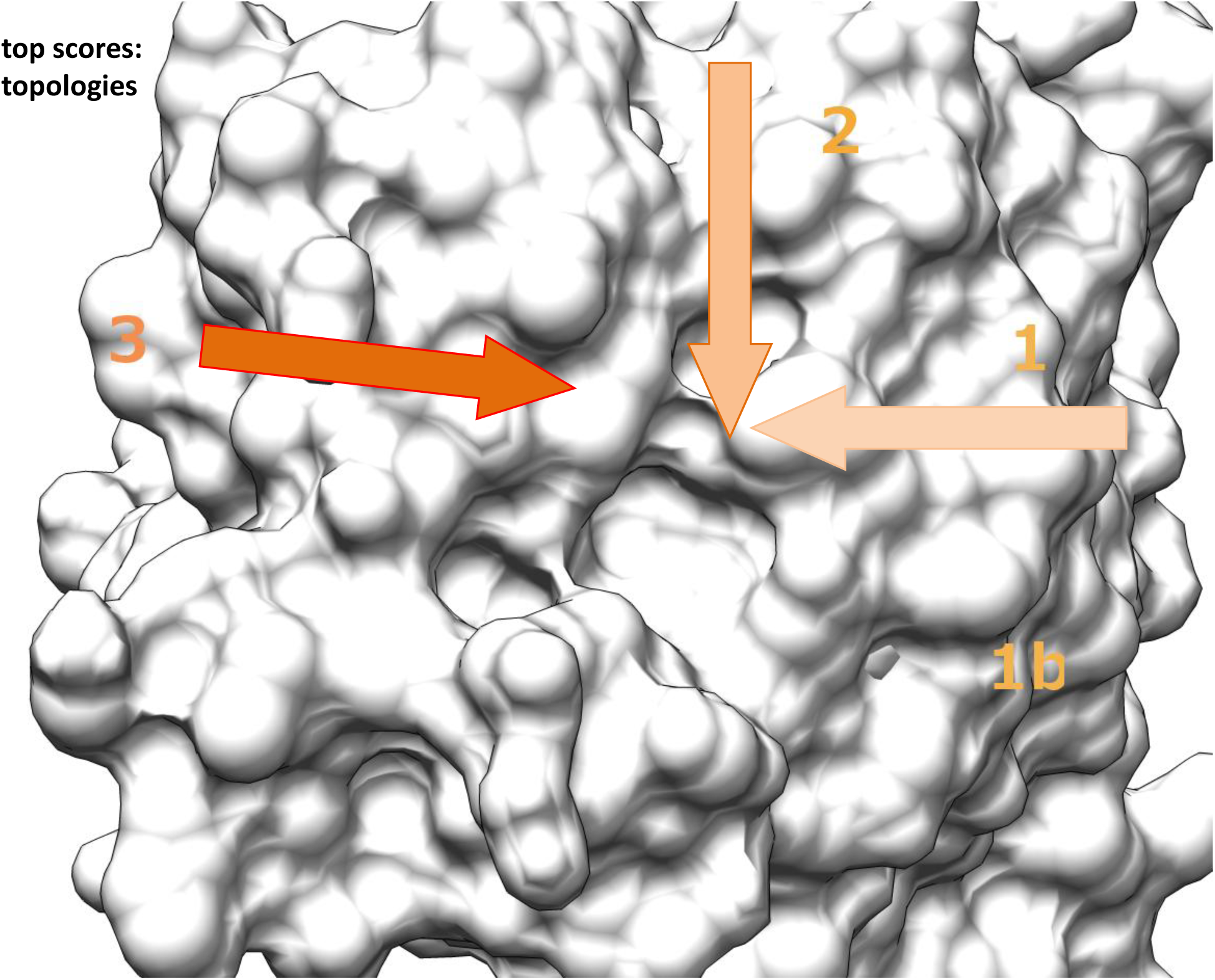
H-bonding of glycan binding and topology of reducing end interaction with VIP36. **A** Comparison of H-bonding. The glycans Carb165 identified in the docking approach were used to establish the binding site geometry with top scoring ligands 9x15. Ligands were evaluated in H-bonding (0.4 Å, 20° relaxed) and the % of bonds formed is graphed. Amino acids are indicated. **B** Topology of the VIP binding site measured by virtual screen. Top 15 glycans of Carb165 were analyzed for binding and found geometries are indicated and labeled with 1-3. The base of the arrow indicates the reducing end of the carbohydrate, the arrowhead the direction of the antennae of N-glycans. Fig. 3 is shown in similar molecular orientation. For comparison also Fig. S2 can be used.

### Contrasting conformers: Ranks and Differences

The subgroup of complex N-glycans scores on average in the first position versus the grouped N-glycans of high mannose type (rank 2), Fucosides (rank 3), Sialosides (rank 4), N-glycan hybrid (rank 5), O-glycans (rank 7) and Other carbohydrates (rank 8) Fig. 5). The plot in Fig. 5 combines the results of all conformers that have been here analyzed.

**Figure 5.**
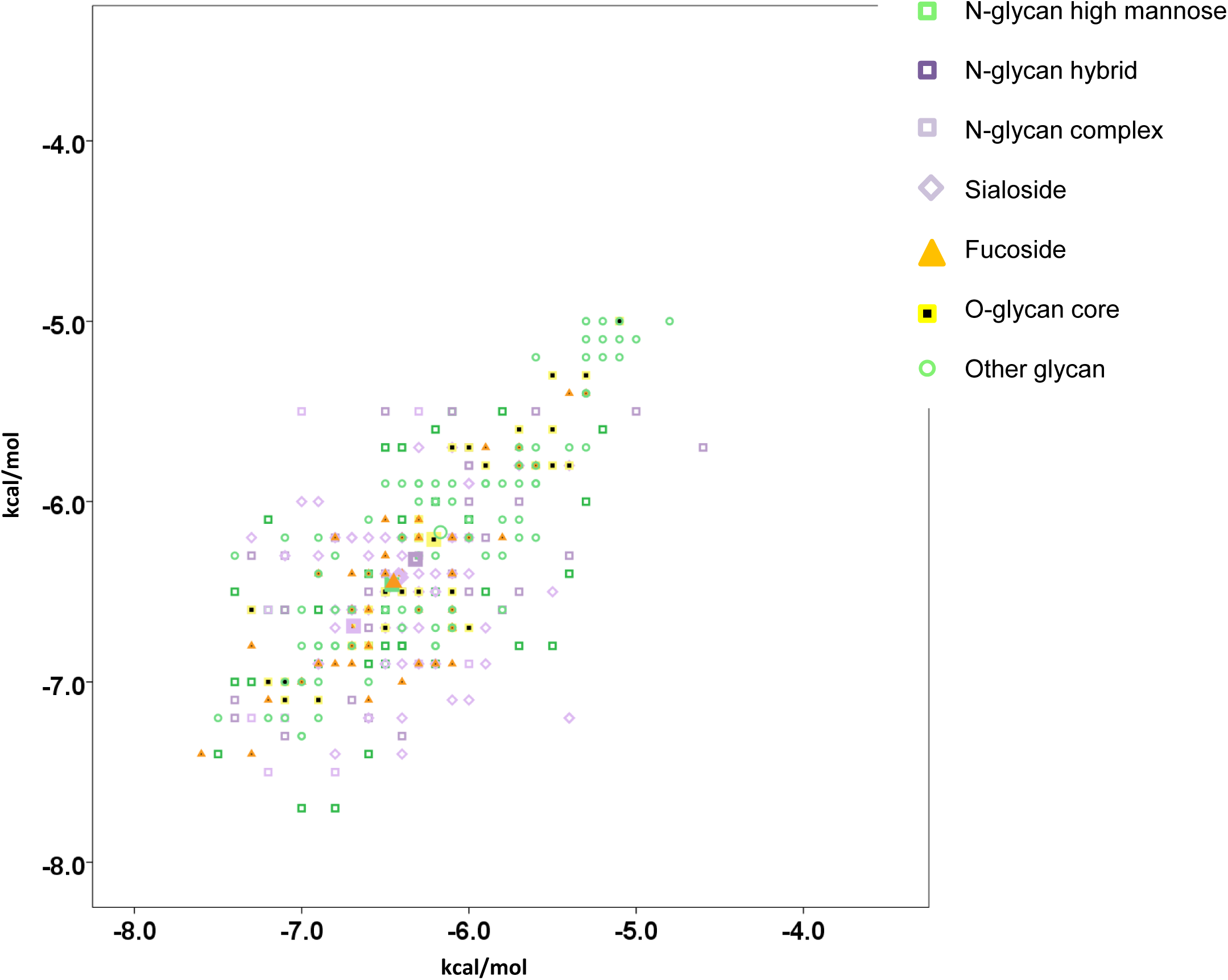
Graphical presentation of Carb165 interactors of VIP36. A Three dimensional plot of all conformers 2E6V, 2DUR-A and 2DUR-B of VIP36 in all-against-all manner (within sub-libraries). Carbohydrates are listed in groups of N-glycan hybrid (violet box), N-glycan high mannose (green box), N-glycan complex (light violet box), Sialoside (light violet diamond), Fucoside (orange triangle) and O-glycan core (yellow square). Other glycans that were not included in this nomenclature are indicated (n=44)(green thin ring). 2DUR-A was graphed against 2DUR-B and 2E6V (each holo). Sialosides include carbohydrates of O-glycan feature if these are sialylated, O-glycans include glycans that are fucosylated. The graph uses scatter plot axes definitions. The runs consisted of 3 sessions with 159 carbohydrates each. The group centers are indicated in large symbols.

Complex N-glycans clearly differ from all other grouped glycans with significantly higher binding affinities (p<0.04)(Table S2A, B), except for the difference of these to N-glycans of high mannose type (p=0.07). Fucoside affinities also differ from some other glycan affinities at significant cutoff (p<0.04), just N-glycans high mannose, Sialosides and N-glycan hybrid fall in the same range of binding strength.

Therefore, among N-glycans significant differences can be established in this docking procedure. Provided the top scoring list prevails in cluster-multiplied approaches, some molecules previously shown by select affinity-chromatography (N-glycans) (Kamiya et al 2005) are also revealed in this simple approach. O-glycans do significantly differ in binding strength relative to N-glycans of complex and high mannose type, Fucosides and Sialosides (p<0.02), which all show higher affinity.

### Topologies of putative glycan interactions

Concerning the interaction of glycans of the Carb165 collection, three different topologies can be discerned. One set of glycans enters the binding cavity from Glu98 described as topology 1 or further 1b (7 / 15). Another group of Carb165 glycans approaches the binding site from His190 in topology 2 (2 /15)(Fig. 4B). Topology 3 shows the carbohydrate approaching the cavity from Tyr164 towards the centre of the binding cavity, towards the bridge in between Asp131 and Gly260 (6 / 15).

Whereas the topology seems rather simple, some O-glycans, and possibly many other low scoring Carb165 residue models, may also contact the binding site from within the central area projection with the reducing end GalNAc residue being elongated towards the centre and multiple indentations.

### Select glycan interactors

Considering the interactions listed in Fig. S2 a number of interesting features of bound carbohydrates should be mentioned. The preferential interaction with N-glycans of high-mannose type is restricted to intermediates of the Golgi-Mannosidase (Golgi-Man) I, an enzyme which is localized to the cis-Golgi (Stanley et al 2009), and is not prevalent for Glc-containing high-mannose precursors. The latter have been implicated in protein folding and quality control (Hebert et al 1995). The final product of a fully trimmed and shortened high-mannose glycan, Man6-[Man3]-Man6-[Man3]-Man4-GlcNAc4-GlcNAc (9; Fig. S3A) binds to VIP36 with distinctly lower energy. The product of endomannosidase, that may function as a failsafe device to cleave residual Glc-Man residues of proteins in the Golgi (Roth 2002), interacts with VIP36 with some intermediate affinity (6). Also the product of α—mannosidase III binds with intermediate affinity, and is itself a substrate for lysosomal degradation (8). Alpha-mannosidase III is thus far described as an alternate backup in most cell types in mice if Golgi-ManII fails (Chui et al 1997; Moremen 2002). Yet, α-mannosidase III cannot fully replace Golgi-ManII since animals develop immune complex diseases in Golgi-ManII null mice (Haltiwanger and Lowe 2004).

A surprising finding is the high affinity of a Golgi Mannosidase II glycan (Fig. S3B) product (22) as well as its substrate (21) to VIP36. Elongated hybrid glycan chains also interact that had not previously been trimmed, although at lower affinity. However, since both, the substrate and product interact at high affinity, the reaction may have reached a “watershed” point of sorting where modified and not yet fully glycanized proteins are subjected to bimodal sorting (Fiedler et al 1996) and bound for backward traffic or the forward route of intra-Golgi transport. How galectin interactions would affect sorting decisions in the Golgi apparatus at this transport stage remains speculative (but see below). These may be stimulated by further GlcNAcT-V (Mgat5) activity in the Golgi cisternae which may enhance glycoprotein binding.

Complex glycan preferences are also considerable with the GlcNAcT-IV product being of second highest energy of interaction (38; Fig. S3C)(kcal/mol). Simple, biantennary complex N-glycans have high but complex tetraantennary products with a bisecting GlcNAc introduced by GlcNAcT-III have lowest energies of binding. If the VIP36 receptor had a sorting role, fucosyltransferase VIII (Fut8) and GlcNAcT-IV substrate turnover may be preferential for generating the ligands for inclusion in exocytic carrier vesicles / tubules. Comparing the top row of glycan affinities (Fig. S3C) it can be seen, that the top interacting molecule displays branching at the Man3 residue (38) but not Man6 (41) and is fucosylated unlike the third ligand (44). Interestingly, LRP-1 (LDL-receptor related protein-1) endocytosis and secretion into the bile may implicate core fucosyl modification as a sorting signal in both, the endocytic and exocytic pathway (Lee et al 2006; Nakagawa et al 2006). In a separate test using the identical docking parameters it was consistently found, that the top scoring complex N-glycan (β-D-GlcpNAc-(1-2)-α-D-Manp-(1-6)-[β-D-GlcpNAc-(1-2)-[β-D-GlcpNAc-(1-4)]-α-D-Manp-(1-3)]-β-D-Manp-(1-4)-β-D-GlcpNAc-(1-4)-[α-D-Fuc-(1-6)]-β-D-GlcpNAc) (38) with or without the core Fuc in α1-6 linkage was scoring at least −0.7 kcal/mol better with the fucose residue added.

The terminal end preferences are displayed in Fig. S3D. Not all enzymes that lead to processing modifications are known. The lactosamine motif seems of preferential energy of interaction, yet, competitive binding or bivalent interaction of ligands with multiple VIP36s seems the sole structurally plausible solution since the N-glycan core is of equal / higher affinity (see above). Comparing the terminal glycan preferences it is surprisingly found that residues β-D-GlcpNAc-(1-3)-β-D-Galp-(1-4)-β-D-GlcpNAc (VI; Fig. S1) show the overall highest affinity of only three linked monosaccharides, unlike the low scoring trimannose of e.g. 2E6V or bimannose of 2DUR-A each scoring with −6.2 kcal/mol. The score allocates rank 4 to the former trisaccharide in global numbering (VI; Fig. S1). In terminal ends, the fucose residues in α1-2 linkage do not increase the binding energy as evident in comparing mono-fucosyls present or absent on trisaccharides (51/56, 53/54; Fig. S3D). This indicates that a single fucose residue as such does not elevate the binding energy in this virtual screen. Furthermore, in N-glycans containing aforementioned trimannose residues, sugars surrounding these will contribute equal to or more than 20 % to the binding energy.

### Glycan interactions in specific and “average” conformers

Since different conformers have been found in the unit cell and in different experiments of crystallization, a subset of these were here compared and summarized. Only single runs of complete libraries were analyzed (without the structure test). Comparing the 3 conformers 2E6V, 2DUR-A and 2DUR-B, the top 15 Carb165 glycans display little variability on average. Summing all 3 rank 3 is filled with a N-linked glycan of high mannose type and by a complex fucosylated N-linked glycan, is preceded by a glycan with the four motif characteristics Lewis C, Lactosamine, Neo-Lactosamine, GSL Lacto series (rank 2), and a fucosylated glycan containing the lactosamine motif (rank 1).

## Discussion

The purpose of this study was to demonstrate carbohydrate binding of VIP36 and to analyze high-affinity interactors in a large-scale docking with 165 glycans including 9 N-glycan complex, 19 N-glycan high mannose, 17 N-glycan hybrid, 17 O-glycan core, 27 Sialoside, 25 Fucoside and 51 other glycan residues.

### N-glycan interaction

In accordance with the previously discovered homology of VIP36, a cargo receptor, with leguminous plant lectins, the present study shows that the residues Asp131, Asn166 as well as His190 are involved in calcium coordination and provide H-bonds to glycans similar to the D1 arm Manα(1-2)- Manα(1-3) of a Man-Man-Man carbohydrate (Satoh et al 2007). Mannose residues in this library of long sized carbohydrates including branched glycans of N-or O-linked glycan modifications frequently bind to the central binding indentation of VIP36 with mannose residues of the D1 or D2/D3 arms within the Asp131 – Asn166 hydrogen bonding pattern (O-linked glycans did not contain mannose). The only residue that shows top scores of binding energy of the N-linked carbohydrate type is number II (Fig. S1 and S2), a complex carbohydrate that does not contact these residues via hydrogen bonds of their mannose groups but places the GlcNAc residues of the core (II) and terminal branch in the hydrogen-bonding distances. These binding interactions may have previously been overlooked in discoveries using serial lectin affinity chromatography (Kamiya et al 2005). In these studies a M8 high mannose ligand, a product of ER mannosidase II was binding to the lectin with highest affinity, in the present study the ligands with very high mannose substitution show less affinity with at least 1.1 kcal/mol energetic difference to the top scoring M7, an intermediate in the Golgi mannosidase Golgi-Man I pathway (Fig. S3A, 10). In a second analysis by lectin affinity chromatography the M8 high mannose residues also showed the highest affinity to VIP36 closely followed by M9 and M7. Whereas the binding of VIP36 was slightly pH-dependent, VIPL showed remarkable reduced affinity at lower pH and preferably bound to M9 (Kamiya et al 2008), an early intermediate. In the present study, the subsequent pathway intermediate M7 interacted with 1.6 kcal/mol higher affinity with VIP36 than the early form M9.

Consistent with the previous analysis, in this unbiased screening I also found the low affinity interactions of the monoglucosylated M9. Moreover, two-fold glucosylated high mannose glycans, substrates of endomannosidase, display less interaction with VIP36 than M9. Quality control mechanisms proposed for VIP36 in the early secretory pathway would here be unconfirmed and likely be unrelated to glucosyl-groups to maintain a transient regulatory cycle of protein folding quality control. This is proposed for the CNX/CRT ER proteins and their glyco-ligands in the secretory pathway (Hebert et al 1996; Ellgaard et al 1999; Hebert et al 2014). While this glucosylation pathway seems excluded for VIP36 based on carbohydrate affinity, a dual function of VIP36 could include the “watershed" point of sorting with both, the Golgi Mannosidase II substrate/intermediate and Golgi Mannosidase II product of equal affinity (Fig. S3B). These two carbohydrates contain one GlcNAc residue each and the VIP36 lectin affinity may be used to halt the secretion of transmembrane / secreted forms of glycoproteins in the medial Golgi up until further modifications ensue. These modifier reactions could install the second GlcNAc group on the α1-6 mannose branch by GlcNAcT-II and could then allow for transport and processing in the Golgi. This mechanism would ensure that accessible N-glycans become more highly glycoslyated and are converted to complex-type glycan forms. A number of other scenarios could be envisioned using fucosyl modifications and further addition of carbohydrate termini as modifiers of the bulk transport in quality control. It is expected, that later extended modifications (Stanley and Cummings 2009) are primarily added to the α1-6 branch of the N-glycan since α1-3 branches shows less mobility as demonstrated in the recent RMSD analysis (Jo et al 2013). The proper installed GlcNAc on the α1-6 branch would then, once further processed from bi-antennary to tri-antennary preferentially on the other branch (see below), once more bind to the VIP36 and be transported or apically sorted since this complex glycan displays highest affinity (Fig. S1).

Another interesting family of proteins are the p24 proteins (Fiedler et al 1996; Fiedler and Rothman 1997). These, also known as EMP24/GP25L/Erp, are another class of cargo-receptors that have been described in COPI-coated vesicles and in the Golgi complex, and newly described functions include cargo clustering and recognition of glycosylphosphatidylinositol (GPI)-anchored proteins (Theiler et al 2014). Thus multiple lectins are likely found in the secretory pathway that each may regulate inclusion into transport vesicles and mediate sorting. In *Drosophila* Wnt proteins require Evenness interrupted (Evi/Wls) for transfer to the plasma membrane while at the same time p24 proteins are necessary for anterograde traffic – glycosylation is furthermore shown to be the targeting signal in apical transport: modifications of Wnt11 include complex N-glycosylation, it is secreted apically whereas Wnt3a which is only high-mannose modified is transferred to the basolateral cell surface in MDCK cells (Buechling et al 2011; Yamamoto et al 2013). Wnt3/5 in *Drosophila* is also N-glycosylated, yet, the role of its glycan modification in the early secretory pathway has not been clarified.

### Glycosphingolipids in development

The third scoring glycan included the GSL ganglioside GM1 in this screen. Assuming that glycosphingolipids are involved in apical delivery as originally envisioned (Holthuis et al 2001), it is tempting to speculate that this interaction is causally related to raft sorting at the exit stations of the Golgi apparatus. The benign effects of mouse β1-4-GalNAc-Transferase knockouts with abolished GM1 synthesis, that only showed effects late in mouse life would have to be explained in view of high affinity GM1 interaction. If sorting platforms would use these together with VIP36 as adaptors, the mouse would alternate these with other raft GSLs. In this model of hexosaminidase deficient and specific knockout mice, abundant GSL GM3 and GD3 were found, but no ganglioside GM1 production (Proia 2003; Furukawa et al 2004). Advances in subcellular localization of glycosphingolipid biosynthetic reactions have pin-pointed to the trans-Golgi for ganglioside and globoside synthetic machinery, and vesicular intra-Golgi transport been ascribed to the former but not to the latter: the TGN was ascribed a role for FAPP2-mediated glucosylceramide (GlcCer) import which is the substrate for specific globoside biosynthesis (Gb3 synthase etc.) and lactosylceramide (LacCer) is synthesized earlier in the Golgi complex if channeled into the ganglioside pathway (D’Angelo et al 2013).

It is rather unlikely that globosides may replace gangliosides in VIP36 binding since globosides are not essential for mouse development and e.g. neither is GM3 - in a VIP36 or raft-based model of sorting this may be proposed. Yet, infants bearing a point mutation in the GM3 synthase gene develop an early-onset epilepsy syndrome (Jennemann and Groene 2013). Most importantly, it has been shown that the Lc3 synthase (β1-3-GlcNAcT-V) gene is developmentally lethal in pre-implantation in mice (Biellmann et al 2008), presumably affecting events as early as adhesion of cells (Hakomori 2008). This would suggest the pivotal role of lacto-series / neo-lacto series of glycosphingolipids. Later analysis showed no essential requirement of the Lc3 synthase in transgenic mice deletion, since mice survived to term (Kuan et al 2010). It is possible that in these three studies (see Review (Jennemann and Groene 2013)) a switch from lacto-to globo-and ganglio-series of GSLs had impact on the cellular growth and differentiation or strain differences contributed to the different results.

Moreover, it is possible that complications affect interpretation of genetic differences and variation possibly due to the different contribution of N-linked glycan chains and their modification since these or both, glycans of lipids and proteins, are used as primarily sorted products. Thus, could VIP36 provide the adaptor for linking the “sorted glycan” and the sorting coat? The raft model includes the sorting of proteins and these may attach to lipids as co-sorted cargo. On the other hand, a raft may be distinguished from other membrane areas and sorted via receptors recognizing a lipid attaching to VIP36, which as sorting platform gathers co-sorted proteins among other cargo. Whether, in this model, VIP36 could provide a sink for 2 sorted ligand classes collectively segregated from other molecules remains to be studied. Since the same enzyme likely adds GlcNAc residues to nascent lacto- and neo-lacto series of GSL and to N-glycan chains it is difficult to predict the outcome of above described experiments with respect to sorting and availability of ligand. Furthermore, of interest are recent findings, that in addition to VIP36, galectins among other cargo receptors are involved in clustering and apical transport of glycoproteins (Mishra et al 2010).

The most likely candidate for an essential function of glycosphingolipids and their headgroup is glucosylceramide demonstrated with a gene Ugcg-knockout mouse. These mice are embryonically lethal and display defects in the ectodermal cell layer at E7.5 due to malfunctioning in GlcCer synthase (Yamashita et al 1999). GlcCer synthase expression is widely found and seems enhanced in this cell layer. It has previously been established that GlcCer is differentially sorted in the apical and basal pathways and this provides evidence for the lipid-based sorting machinery (vanIjzendoorn and Hoekstra 1999; Holthuis et al 2001). It is possible that only the combined effect of loss of all GSLs of the GlcCer series have these detrimental outcomes. The role of galactosylceramide (GalCer) synthase has been controversially discussed and studies in Ugta8a knockout mice have found survival for 90 days with severe demyelination of the central nervous system or death at 25-30 days after birth. Further into this branch of GalCers, it has been found that sulfatide formation by the sulfotransferase (Gal3st1) will also affect the nervous system in myelination as well as male fertility although later and less severe (Honke et al 2002).

Current insight would suggest that although there is an exquisite order of gene-regulation in development and sequential up-regulation of specific glycolipid biosynthesis at distinct time points the fundamental nature of the lipid bilayer and its sorting machinery depends on very few carbohydrates that are added to GSLs or functional redundancy may explain most observations that have been made so far.

### Branches of N-glycans and their lectin affinity

One important result from this screen is the lower interaction of GlcNAcT-V (Mgat5)-branched N-linked glycans (Partridge et al 2004; Boscher and Nabi 2013) that show less binding than the top-binding GlcNAcT-IV-branched glycans, corresponding to the D3 versus D1 mannose arm without GlcNAc (see overview Figs. 6, 7). Since previous experiments in animals in tumor growth have pinpointed to the role of these enzymes and products as decisive modifiers of metastasis and focal adhesion signaling, it may be tempting to speculate on the function of carbohydrate branching with respect to conformational freedom, glycoprotein sorting and, as shown, membrane organizing functions (Granovsky et al 2000; Christiansen et al 2014; Stowell et al 2015). The conformational space of singly branched versus doubly or even triply branched polymers is restricted at each branch point and thus their wandering in space is limited to a smaller number of interactions. In this regard the Mgat5 branch is no exception but since the number of neo-or lactosamine groups added preferentially to the Mgat5 terminus is high, the number of interactions cannot be predicted but may change in their selectivity. If the sorting function that is proposed is fulfilled, VIP36 could bind to the GlcNAcT-IV branch of modified glycoproteins but less to the GlcNAcT-V branch of processed glycans and may properly sort the former but less so the latter. In contrast, the focal contact interactions that have been shown to involve the lectin galectin-3 may be stimulated by increased Mgat5 expression, and this may lead to maintaining a longer residence of Mgat5-modified glycans (Partridge et al 2004). Therefore, assembled complexes of epidermal growth factor (EGF) and transforming growth factor (TGF) receptor and galectin-3 may signal longer and promote the invasion of cells stimulating cell motility. The aforementioned role of galectin-3 in nuclear signaling via the β-catenin pathway that was demonstrated in tumor cell lines would furthermore contribute to the epithelialmesenchymal transition (Shimura et al 2004). In tumour sections, galectin-3 is detected with the peroxidase-antiperoxidase reaction (Mac-2) and in contrast to its expression in the cell lines, the decrease of nuclear localization is associated with tumour progression (Lotz et al 1993). In colon carcinoma the interplay with other galectin family members or the stage specific alterations of cell growth, adhesion or attachment may be different.

While complex N-glycans are certainly important for growth since in GlcNAcT-I knockouts mice do not survive to term (Ioffe and Stanley 1994) not all of their functions were appraised. Interestingly, homozygous GlcNAcT-I -/- cells did not participate in growth and the lung epithelial cell layer in the bronchus demonstrating the dependence on complex/hybrid N-glycans in adhesion (Ioffe et al 1996). Yet, recent results suggest that the oligomannose form e.g. of modified E-cadherin produces stronger cell-cell interactions than the complex or bisecting GlcNAc-modified form (Hall et al 2013) and that an explanation of the phenotype of the mice will require more molecular research. In the analysis of Lec1 versus Pro-5 and LEC10B cells it was suggested that the information content of glycans is being utilized in surface expression and organization. With respect to sorting it is likely that in polarized cells the glycocode would be utilized for polarized apical targeting.

**Figure 6.**
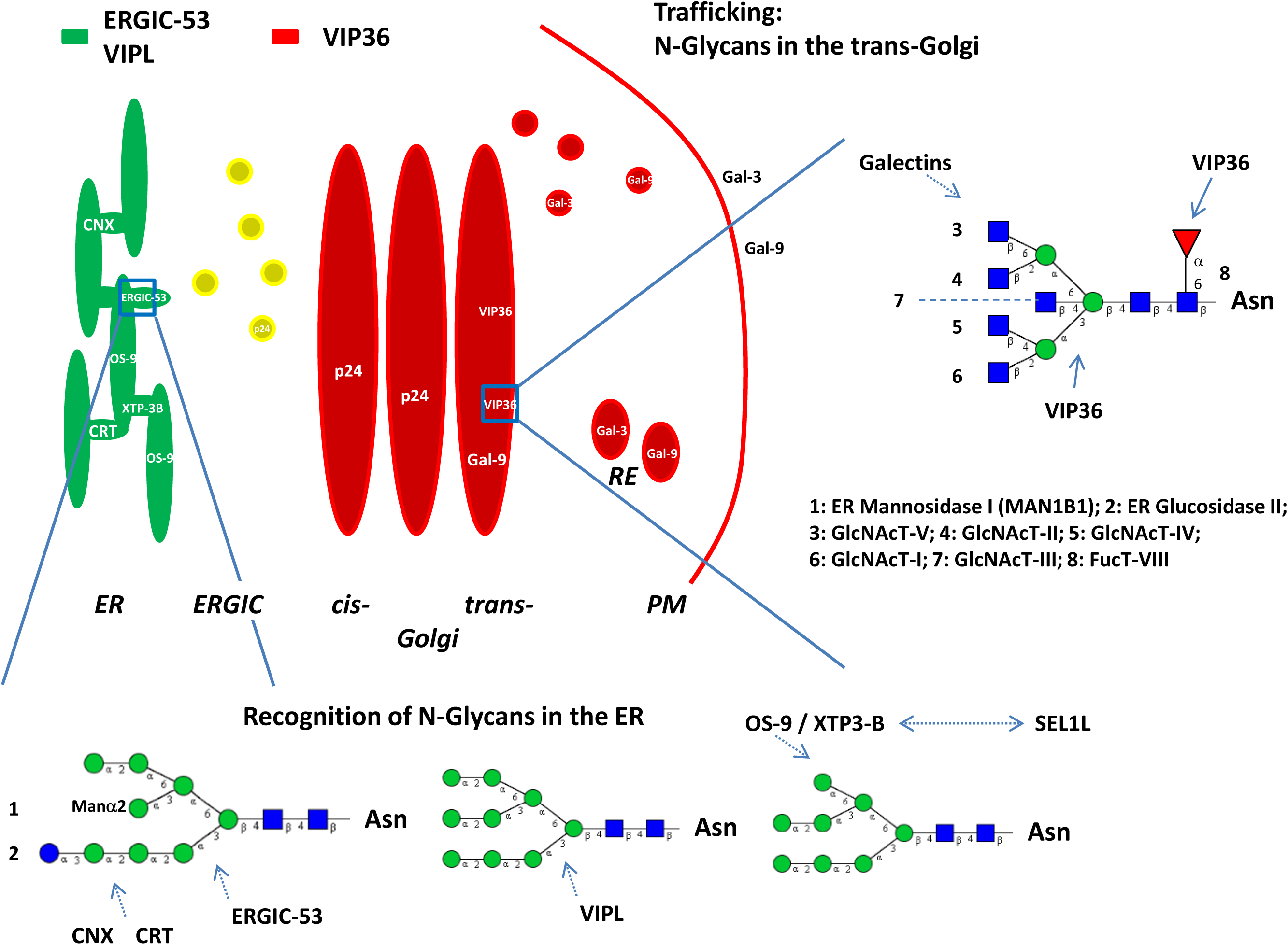
Model of transport and sorting: Overview of N-glycan structure, VIP36 lectin affinity is shown here for the first time for the high mannose and complex type N-glycans. Some enzymes involved in N-glycan processing are indicated. MAN1B1 has recently been identified in the Golgi complex (see (Hebert et al 2014)). ERGIC-53 (green), VIPL (VIP36-like protein), VIP36 (red) and Galectins are indicated and are localized to the ER, ERGIC (yellow; co-localization), Golgi apparatus, RE or PM. The residue labeled Manα2 has not been included in a previous model of ERGIC-53 interaction (Satoh et al 2014) and is indicated for clarity since the M9 (Man9) carbohydrate residue is the described substrate of (1) and (2). Serial lectin affinity chromatography points to monoglucosylated M7 as the high affinity interactor of ERGIC-53. VIPL is described to interact with M9 and M8 high mannose residues with high affinity. Galectin-9 (Gal-9) is a high affinity interactor of polylactosamines. Forssman antigen glycosphingolipid (GSL) was found to interact as well, the exact GSL that binds to galectin-9 in polarized sorting in humans remains to be determined (Svensson et al 2012). Galectin-3 (Gal-3) was found to be involved in a non-raft pathway in apical sorting. VIP36 is modeled to bind to docked core residues of N-glycans involving a fucose residue (1-6 linked), mannoses and N-acetylglucosamines. Preferential interaction of VIP36 with the mannose (1-3) branch makes additional clustering interactions likely for the mannose (1-6) branch. Apical sorting signals would involve both branches of N-glycans and GSLs. No preferential binding site of core fucose 1-6 of all complex N-glycans (Fig. S3C; 38, 39, 41) with VIP36 could as yet be demonstrated. Calnexin (CNX) or Calreticulin (CRT) and lectin ERAD (ER-associated protein degradation) receptors OS-9 or XTP3-B are also indicated. OS-9 binds to M8 - M5 high-mannose N-glycans, XTP3-B binding may be similar (Hebert et al 2014). Both proteins may also directly interact with the membrane protein SEL1L of the ERAD complex via its own glycans. Further information on ER quality control of protein folding and degradation is found in the cited review. The p24 proteins are introduced to mention their interaction with the glycosylphosphatidylinositol (GPI)-anchor of GPI-anchored proteins (p24γ) likely, indirectly or directly, through their membrane proximal α-helical domain (Theiler et al 2014). Abbreviations: ER (endoplasmic reticulum), ERGIC (ER-Golgi intermediate compartment), PM (plasma membrane), RE (recycling endosome).

**Figure 7.**
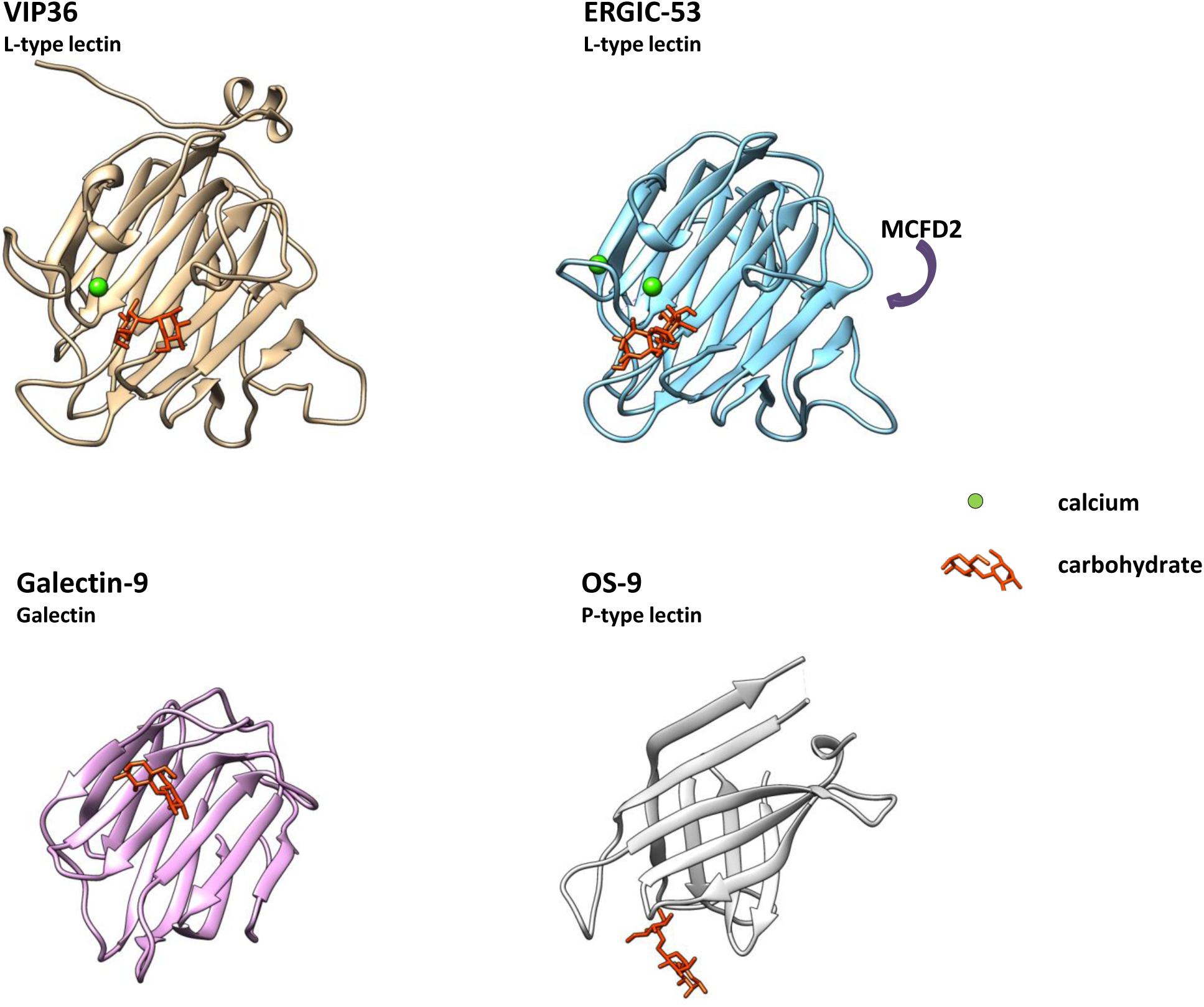
Structural comparison of L-type lectin, Galectin and P-type lectin. VIP36 (2DUR-A; residues 51-301), ERGIC-53 (4GKX-A; residues 31-270) and Galectin-9 (3LSE-A; residues 6-148) were analyzed and aligned in three dimensions with Chimera. P-type lectin OS-9 (3AIH-A; residues 109-228) is shown in similar aspect for comparison. Carbohydrates are indicated and shown in the binding pocket (orange) if present. Calcium is shown in green, top-scoring entries of the similarity search (Table 1) were aligned. Galectin-9 binds carbohydrates in similar position to e.g. galectins 3 and 4. The P-type lectin OS-9 and OS-9like proteins recognize carbohydrates primarily via strand residues, OS-9 binding involves a Trp-Trp motif. Further P-type lectins/mannose 6-phosphate receptor (MPR) homology domain-containing lectins (including the cation-independent and cation-dependent MPR) have been omitted. The glycan binding site of VIP36 and galectin-9 is non-similar, whereas primarily loop residues Ser96, Asp131, Asn166, Gly260, Asp261, Leu262 bind to the carbohydrate in VIP36 it is strand residues in galectin-9 that interact including His61, Asn63, Arg65, Asn75 and Arg87 that do not align. Resolved carbohydrates in the structures corresponded to α-D-man-α-D-man (VIP36), α-D-man-α-D-man (ERGIC-53), β-lactose (Galectin-9) and β-D-man-α-D-man-α-D-man (OS-9). ERGIC-53 has been crystallized in the presence of the recruitment protein MCFD2 (Multiple Coagulation factor Deficiency protein 2)(Nishio et al 2010). The binding face is indicated. MCFD2 binds coagulation factors V and VIII and in an autosomal recessive disorder in mutants causes reduced serum levels. Independent ERGIC-53 binding of glycoprotein ligands without MCFD2 has also been shown

## Conclusions

With respect to the sorting signals it is predicted that the core α1-6 linked fucose would enhance apical transport of N-glycosylated proteins once bound to the sorting machinery. Interestingly, it has been found that in hepatocytes this core fucose may be involved in apical transport (Lee et al 2006; Nakagawa et al 2006). Although speculative, it is possible that the core fucose modification is essential, but loss-adaptation can upregulate GlcNAc bisection by GlcNAcT-III (Wang et al 2005; Kurimoto et al 2014) that in glycan-receptor interaction would exert a similar function as the added core fucose.

The GlcNAcT-V products of N-linked glycans could be anchored to the cytoplasmic coat via soluble clustering receptors and ensuing cargo aggregation which may function akin to coincidence detection on the cytoplasmic face, including cytoplasmic domains of cargo-receptors and luminal soluble galectin-9. Since neither galectin-9 nor galectin-3 have previously been shown to preferentially interact with core-fucosylated N-glycans (Hirabayashi et al 2002) VIP36 may carry out its function by directly interacting with the fucose core. Carbohydrates interacting with each family member of L-type lectins *in vivo* remain to be studied in detail in the future.

The role of VIP36 or VIP36-like proteins in phagocytosis (Shirakabe et al 2011; Huang et al 2014) and its lectin function in clearance of *Staphylococcus aureus, Vibrio parahaemolyticus*, and *Aeromonas hydrophila*, or *Campylobacter jejuni* suggested based on the current results, may be studied in a further project.

Large scale screening including carbohydrate headgroups of glycolipids will require supercomputing in particular if multiple lectins and/or conformers are studied. It could be expected that cancer treatment and the understanding of cancer metastasis would benefit from this undertaking.

## Methods

Structural comparisons were carried through with PDBeFold v.2.59 (Krissinel and Henrick 2004)(searched 8. Feb. 2015).

The docking approach was carried through with structures obtained from PDB (http://www.pdb.org) using the 3-dimensional (3D) coordinates published for 2DUR and 2E6V. The programme used for docking was AutoDock VINA (Trott and Olson 2010). The input and output uses PDBQT structure format. Since the coordinates for 2DUR-A provided ambiguous side-chain conformations that showed disorder in the X-ray analyses, the chosen conformers for aforementioned approach consisted of Glu98(A)-Asn183(A)-((Gly184(A))-Ser185(B) and Thr203(A) rotamers. Within the AutoDock implementation of PyRx version 0.8 from Sargis Dallakyan (http://pyrx.scripps.edu) the grid was set as indicated, which reached a minimal resolution of 0.375 Å for all local docking approaches. Only continuous runs of docking for all Carb165 residues in one session were evaluated (unless otherwise indicated in control simulations). The structural criteria chosen were the formation of double H-bonds to Asp131 and Asn166 for some panels as indicated; this negated often cited limitations of top scores to RMSDs of 0,0. The global approach necessitated a slightly lower resolution for the grid (sizes 51×51×40 Å). The programme utilizes OpenBabel (O’Boyle et al 2011) and AutoDock 4/VINA (Forli and Olson 2012) (http://autodock.scripps.edu) and an uff (united force field) for energy minimization, conjugate gradients with 200 steps and a cut-off for energy minimization of 0.1. Partial charges were added to receptors using PyBabel (MGL Tools; http://mgltools.scripps.edu). No limits to torsions were allowed which resulted in a maximum of attempted adjustments. Single CPU time was up to 150 hours for longest/branched ligands.

The carbohydrate collection “Carb165” was downloaded from the laboratory of Prof. Woods (Tschampel et al 2006) (http://www.ccrc.uga.edu) (with multiple conformers), was selected and filtered for uniqueness against a laboratory collection of glycosphingolipid-glycan headgroups (GSL) that was built using the chemical descriptor-library of LIPIDMAPS and downloaded (http://www.lipidmaps.org) or the Glycosciences website (http://www.glycosciences.de).

H-bonding was determined with ViewDock and with tolerances 0.4 Å, 20° (Mills and Dean 1996). Annotation of carbohydrates was obtained from descriptions and species annotations from http://www.glycome-db.org including the MCS search routine (Ranzinger et al 2009). Drawing utilities were from GlycoWorkbench (Ceroni et al 2008). Prosite searches were carried through with prosite.expasy.org (Sigrist et al 2002). Panther search with http://www.pantherdb.org was used for visualizing multiple sequence alignments and phylogenetic tree results (Thomas et al 2003).

Deep View Ver.4.1 (http://www.expasy.org/spdbv) and Chimera (http://www.cgl.ucsf.edu/chimera) were used for further structural analyses. Energy minimization of structures was carried through with a minimum of 100 steps of conjugate gradients applying the amber ff12sp force field (Wang et al 2006; Case et al 2012) using Gasteiger charges (Gasteiger and Marsili 1980). “Holo enzymes” were thus constructed from the apo-liganded structures by converting side chains rotamers within the binding site as well is within the entire structure for advanced tracking of docked ligands. Ligands were analyzed with ViewDock. Hydrogen-bonding files were further processed in Excel (Microsoft) and sqlite data were analyzed using SQLite (Hipp, D. R.).

Further graphics were calculated or presented with SPSS Statistics (IBM) 21 and Excel (Microsoft). T-tests were carried through with post-hoc testing including skewness, Kolmogorov-Smirnov and Levene tests.

## List of abbreviations

CNX: calnexin
CRD: carbohydrate recognition domain
CRT: calreticulin
EGF: epidermal growth factor
ER: endoplasmic reticulum
ERAD: endoplasmic reticulum associated degradation
ERGIC: endoplasmic reticulum-Golgi intermediate compartment
Evi: evenness interrupted
Fuc: fucose
Fut: fucosyltransferase
G: ganglioside (with class indicator)
Gal: galactose
GalCer: galactosylceramide
GalNAc: N-acetylgalactosamine
Gb: globoside
Glc: glucose
GlcCer: glucosylceramide
GlcNAc: N-acetylglucosamine
GlcNAcT-V: GlcNAc-Transferase V (Mgat5)
Golgi-Man: Golgi-mannosidase
GPI-anchor: glycosylphosphatidylinositol-anchor
GSL: glycosphingolipid
Hep3B: hepatocellular carcinoma cells
Lac: lactose
LacCer: lactosylceramide
Lc: lacto-(with numerator for the number of carbohydrates)
LDL: low-density lipoprotein
LEF: lymphoid enhancer binding factor
Man: mannose
MCFD2: multiple coagulation factor deficiency protein 2
MDCK: Madin-Darby Canine Kidney
NeuAc: N-acetylneuraminic acid
NSCLC: nonsmall cell lung cancer
TCF: T-cell transcription factor
TGF: transforming growth factor
Ugcg: UDP-glucose ceramide glucosyltransferase
Ugt: UDP-galactose ceramide galactosyltransferase
VIP36: vesicular-integral membrane protein of 36 kD
Wnt: wingless

## Competing Interests

The authors have declared that no competing interests exist.

## Authors’ Contributions

KF conceived and carried out the study, drafted the manuscript and prepared tables, figures and graphics. The author read and approved the final manuscript.

## Acknowledgements

Dr. Lavinia Alberi, Department of Medicine, University of Fribourg is thanked for critically reading the manuscript. Prof. Kai Simons, MPI MCBG, Dresden is thanked for pointing out the relationship to galectin and sorting.

## Supplementary Figures/Tables

**Figure S1.**
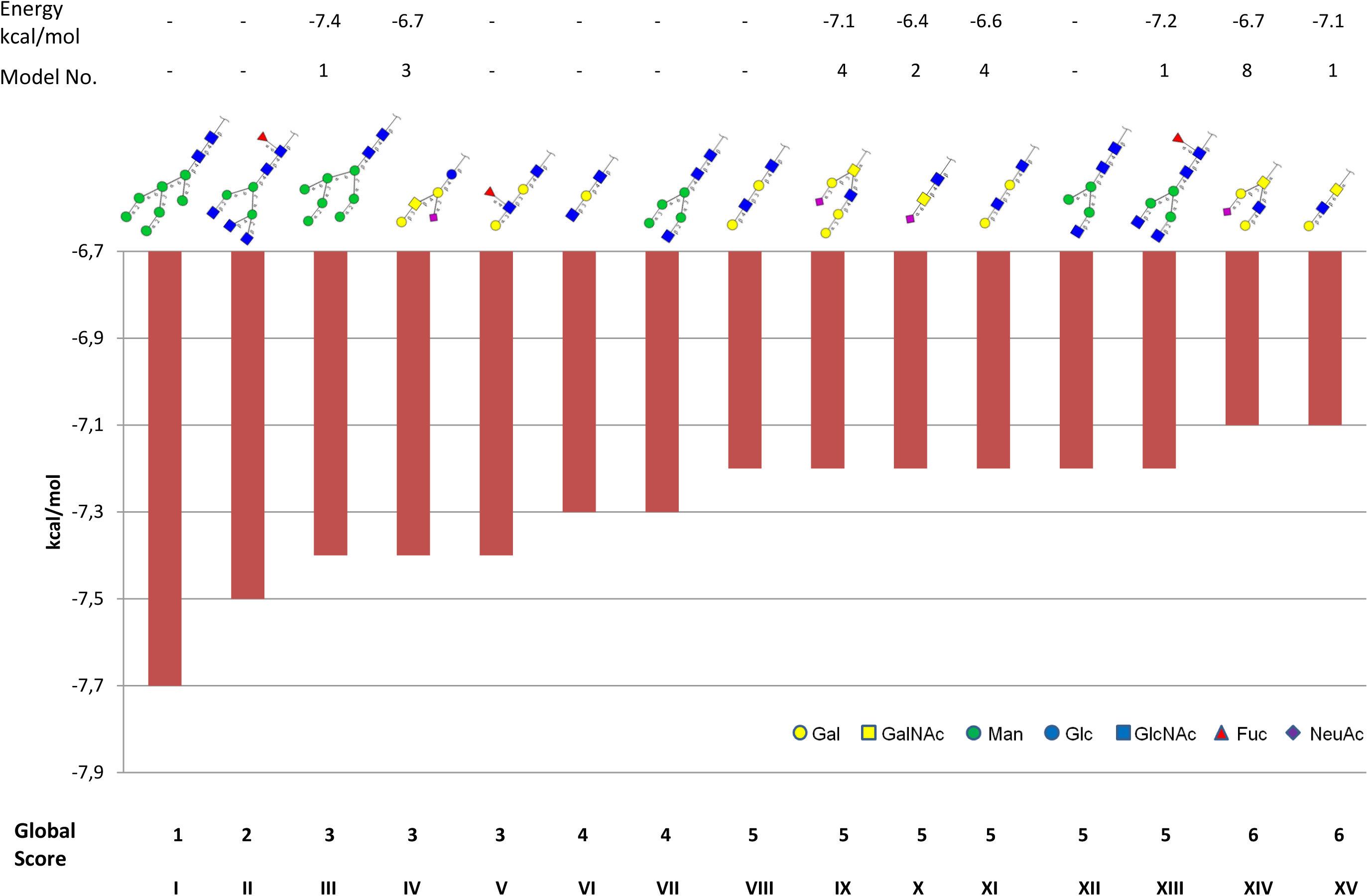
Top Carbohydrates interacting with VIP36. The structurally identified mannosyl ligand of 2DUR-A was removed, the structure was energy minimized and virtually screened. CARB residues interacting with highest binding site affinity are indicated and presented in IUPAC nomenclature. All ligands cover the central binding pocket that provides binding indentations that are seen (geometrically) between Tyr164, Asp261 and Pro 146, Glu 98 covering the central area Asp131, Asn166 (see Fig. 3). Energies indicated above are listed with values for molecules interacting with both, Asp131 and Asn166 via (0.4 Å, 20° relaxed) H-bonding. Model ranking is given for these doubly (or multiply) H-bonded ligands selected. Gobal score refers to ranking which combines CARB and GSL residues neglecting above mentioned structural requirement of forming 2 H-bonds to Asp131, Asn166. Sequential labeling in roman numbers.

**Figure S2.**
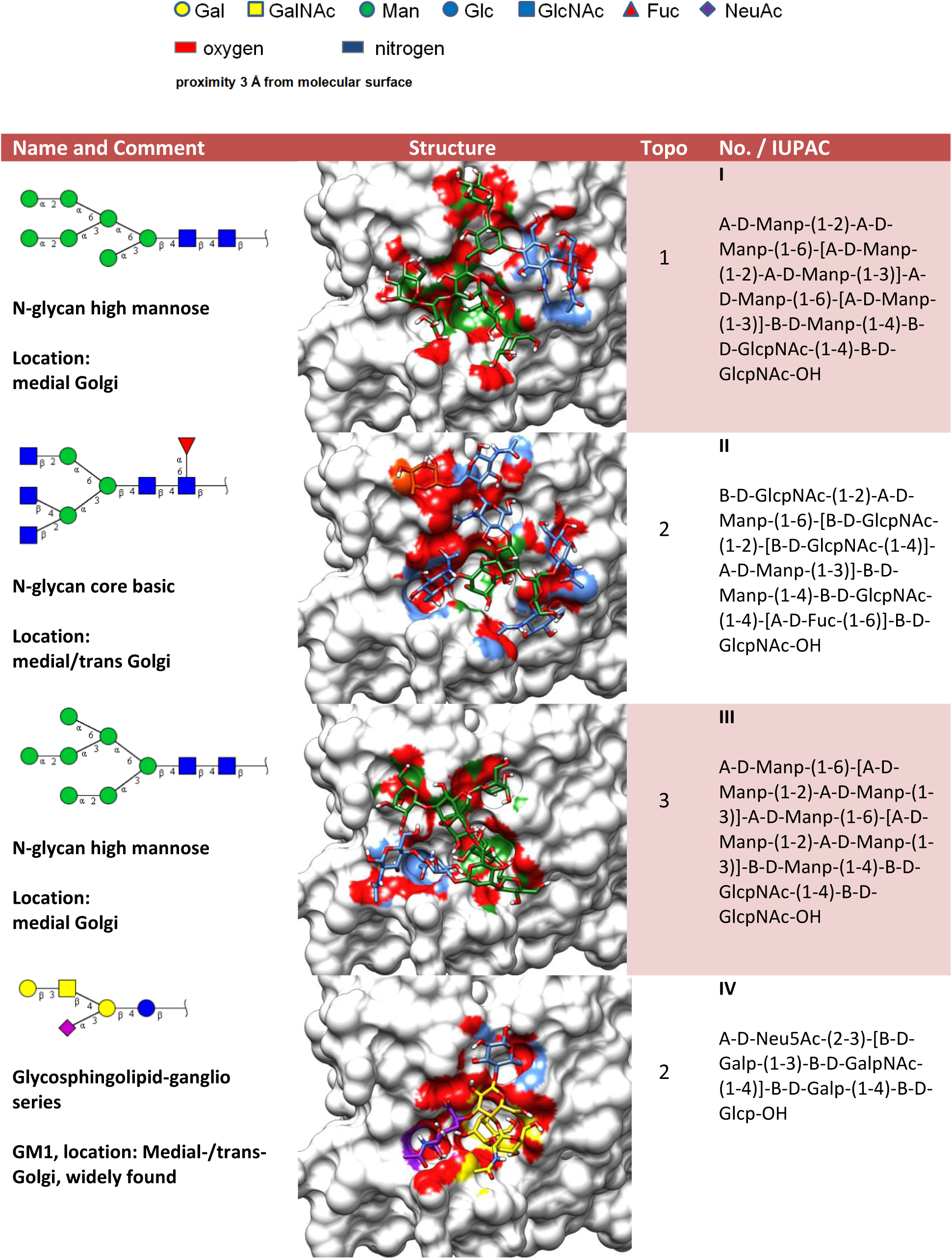
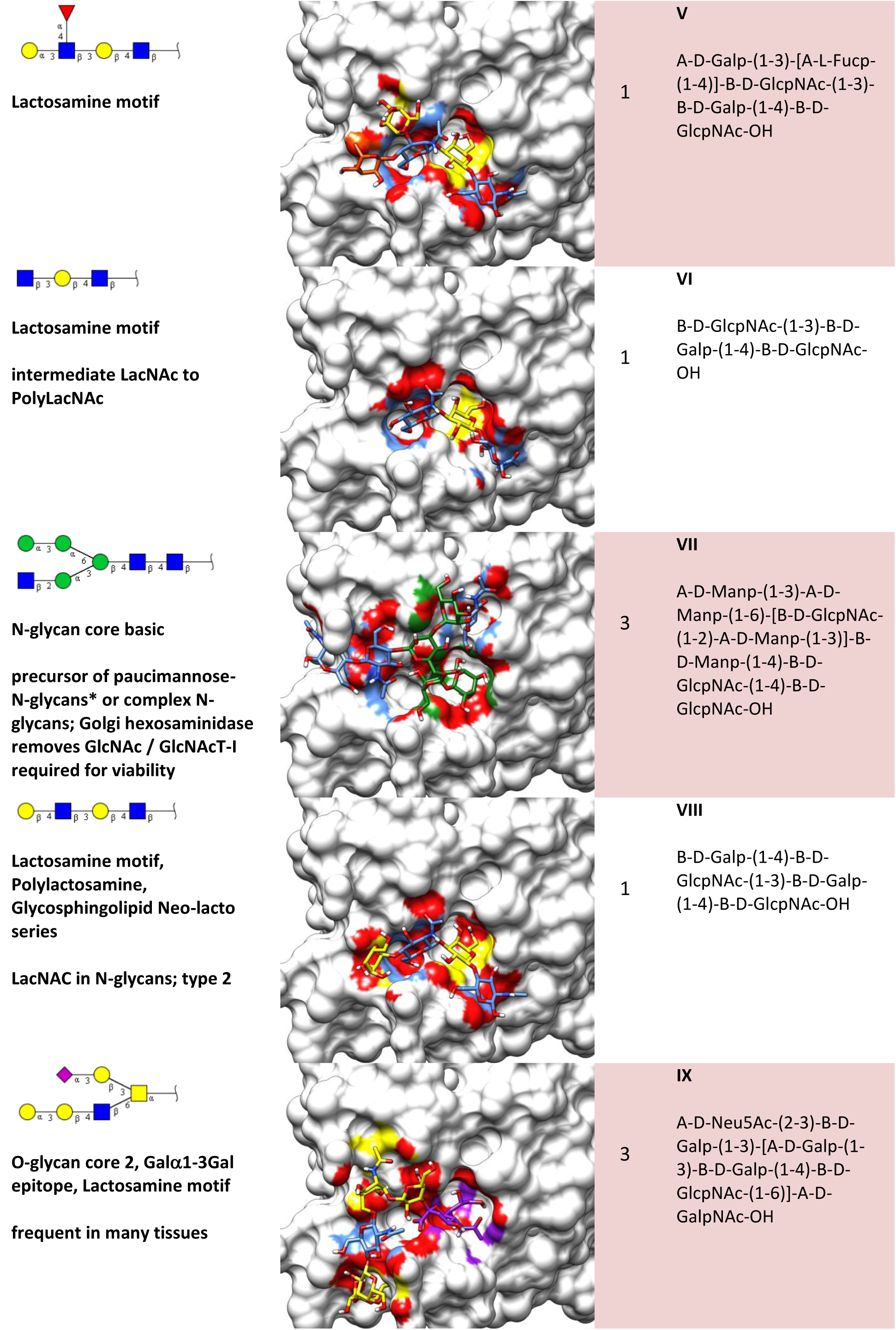
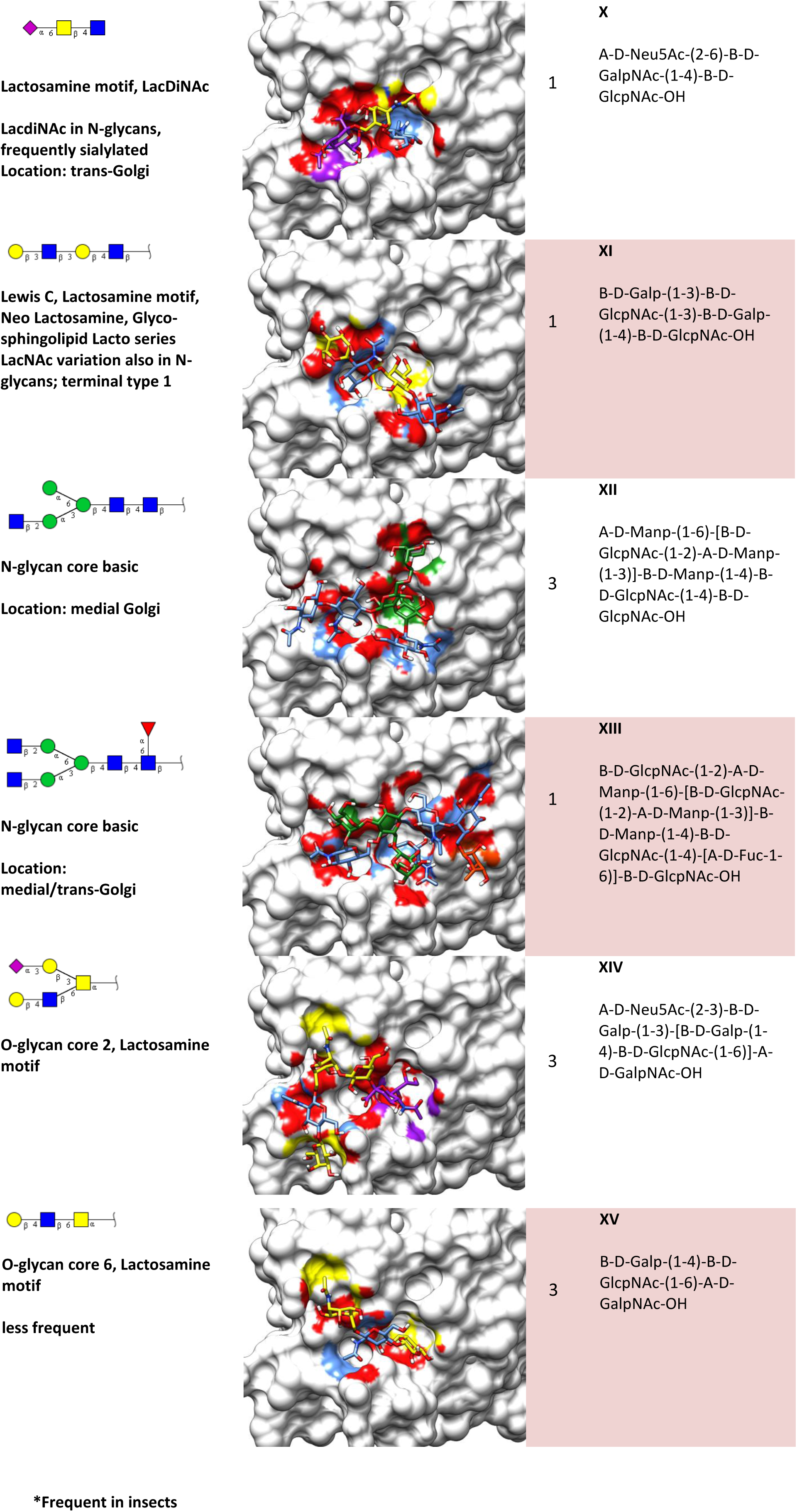
Structure and docking of Carb165-library VIP36 ligands. The ligands identified with highest affinity to VIP36 (2DUR-A) are listed with IUPAC coloring *(Structure)* and imprint onto the VIP36. Heteroatom coloring was utilized to show potential H-bonding interactions. The topography *(Topo)* of the ligand binding reveals orientations of headgroup and cores that corresponded to three axes numbered 1-3 (see Fig. 4B). IUPAC formulas *(IUPAC)* and sequential labeling in roman numbers are indicated.

**Figure S3.**
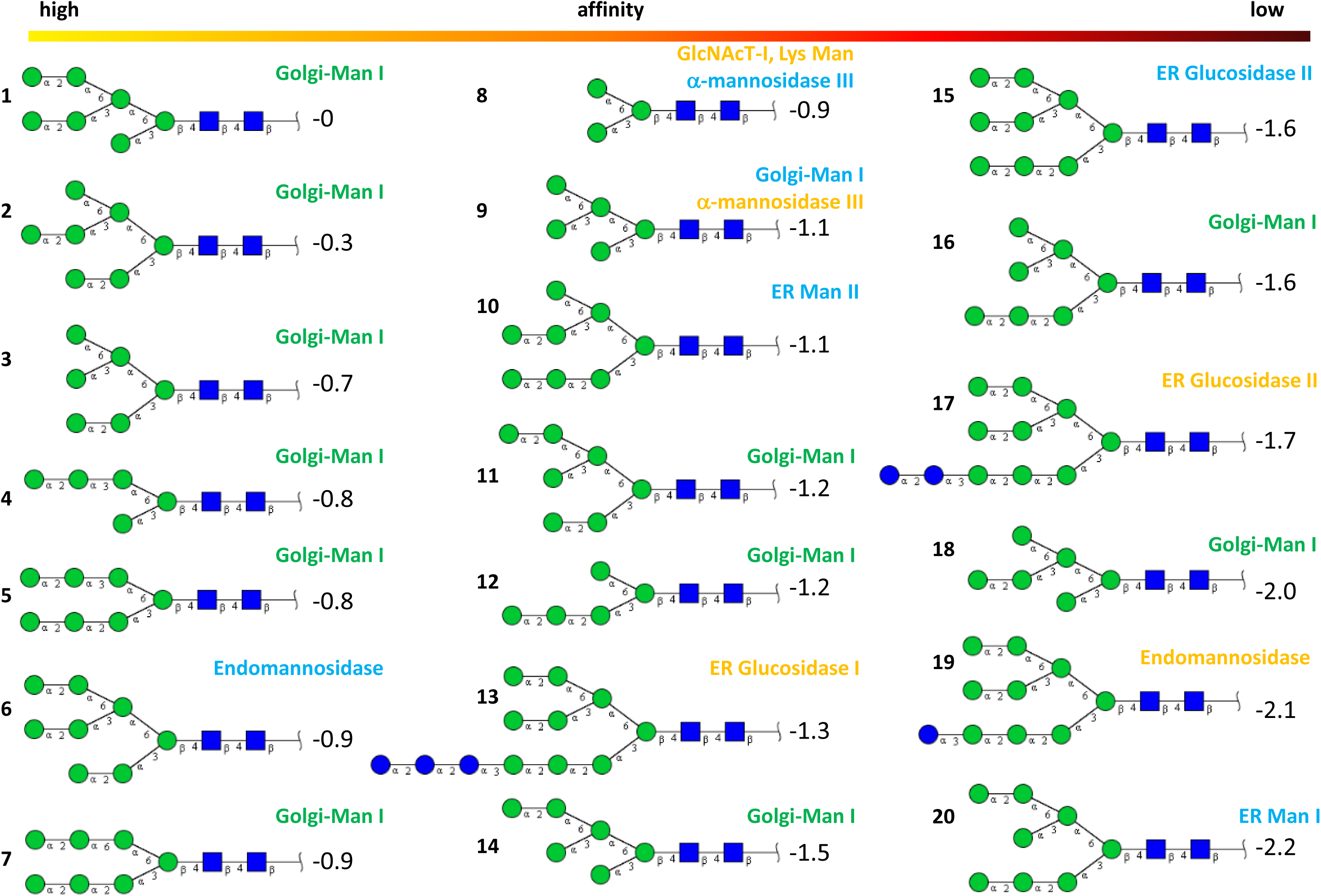
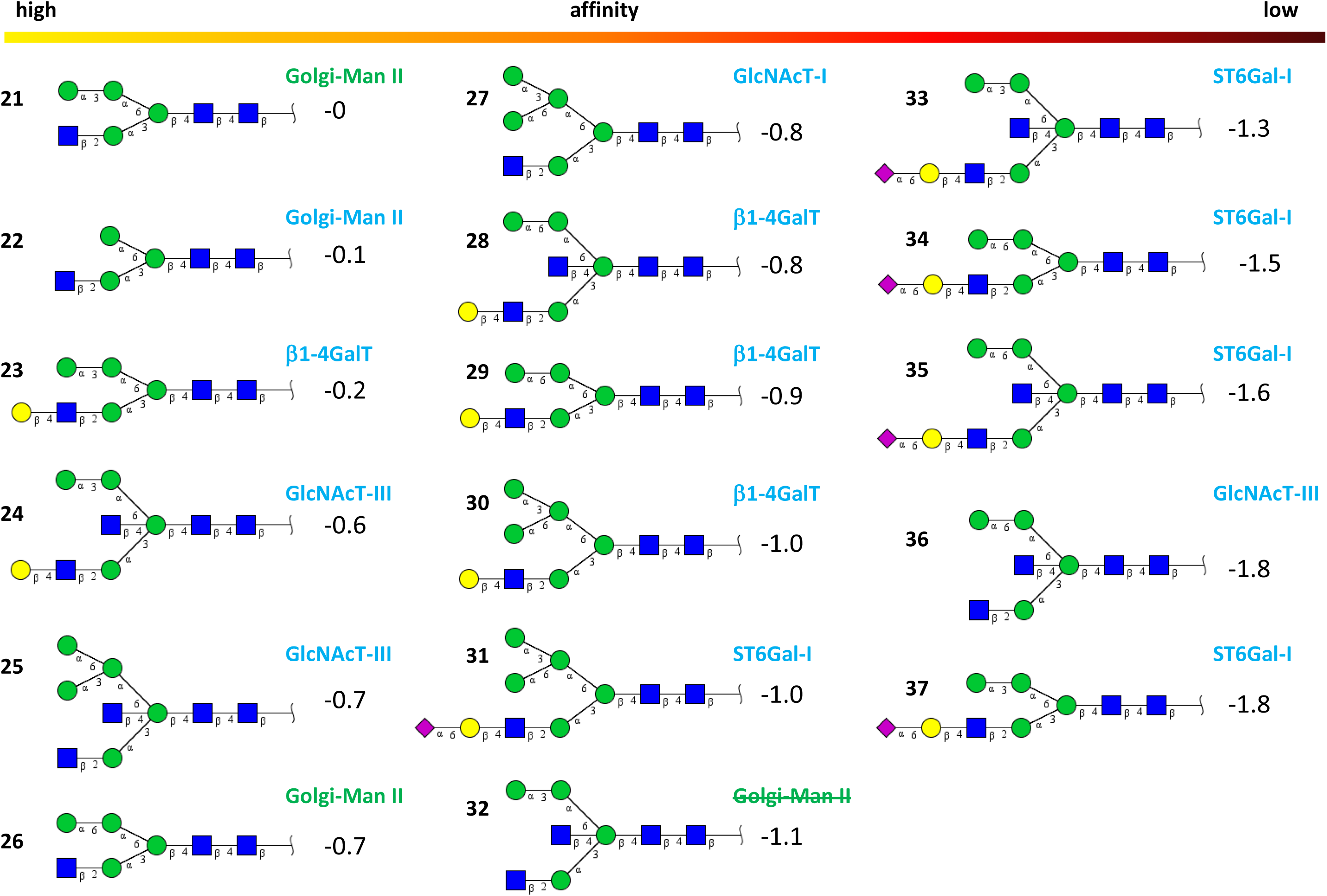
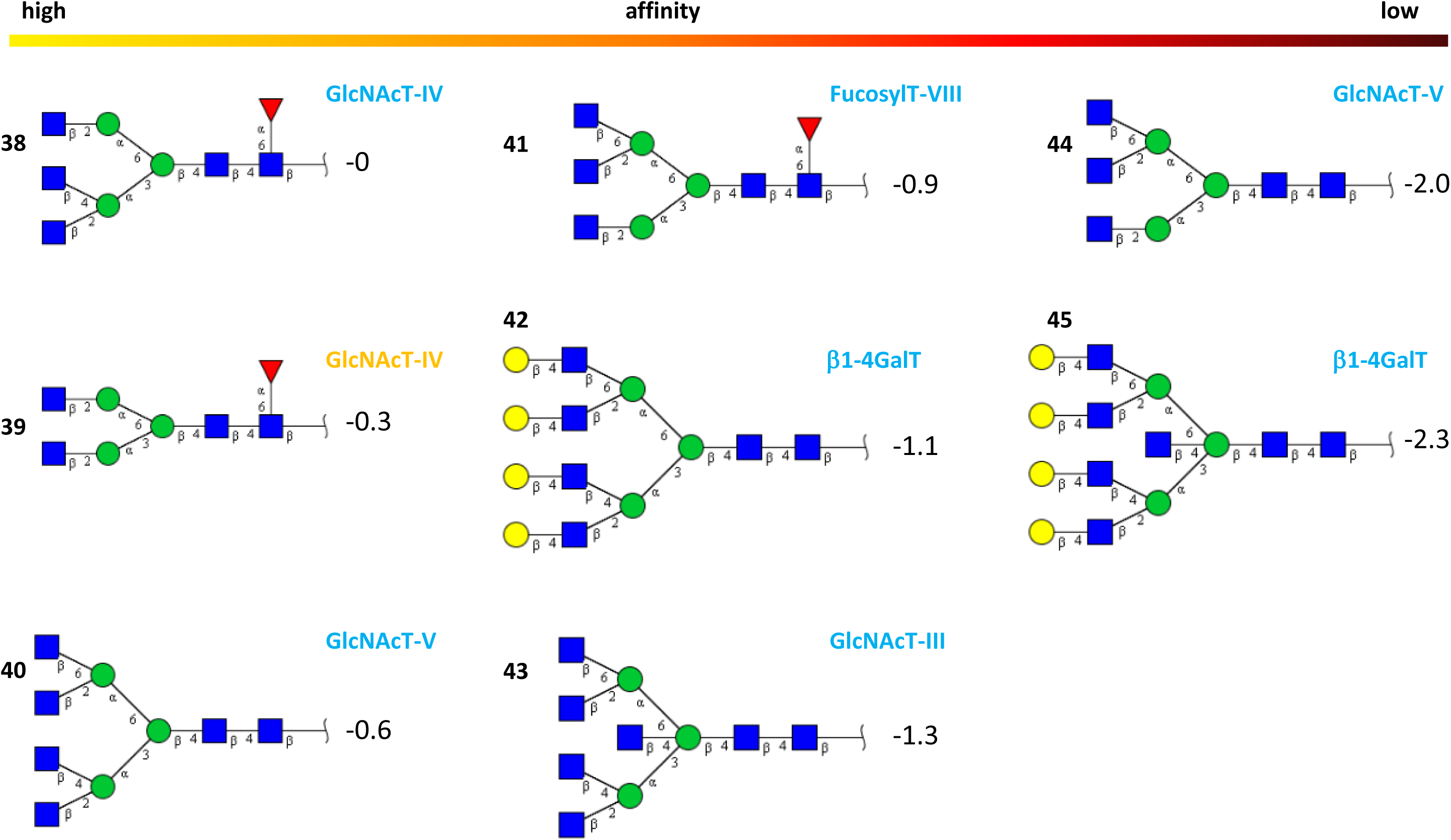
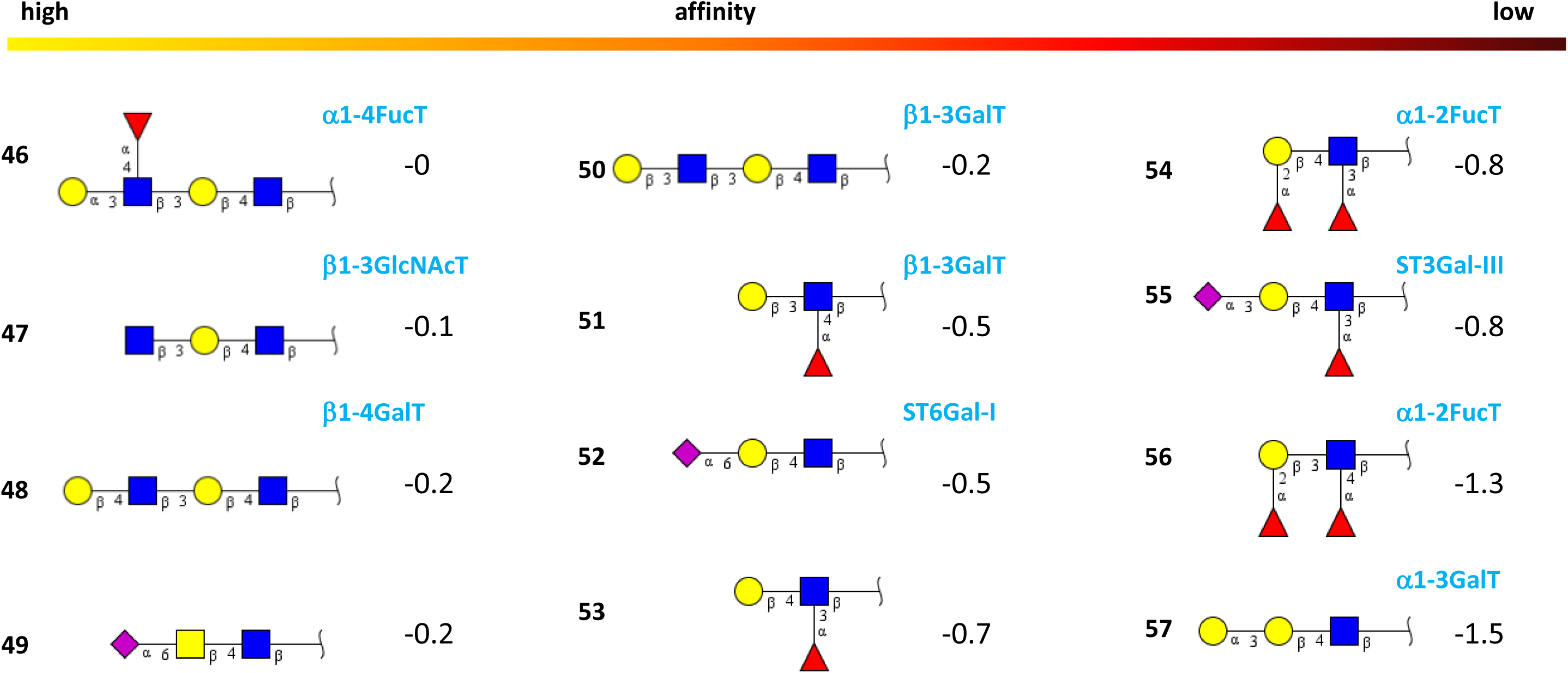
Glycan preference. The carbohydrate interaction of VIP36 (2DUR-A) was classified according to the energy (kcal/mol) and top scoring of each glycan group; Δ values are indicated. **A** High-mannose glycan preference. **B** Hybrid glycan preference. Standard enzyme product and substrates are indicated. **C** Glycan complex preference. **D** Glycan terminal end preferences. Some enzymes with (reference to the text) that are involved in the reaction paths are indicated with edduct (orange), product (blue) and intermediate (green). Sequential labeling is indicated in numbers.

**Table S1.**
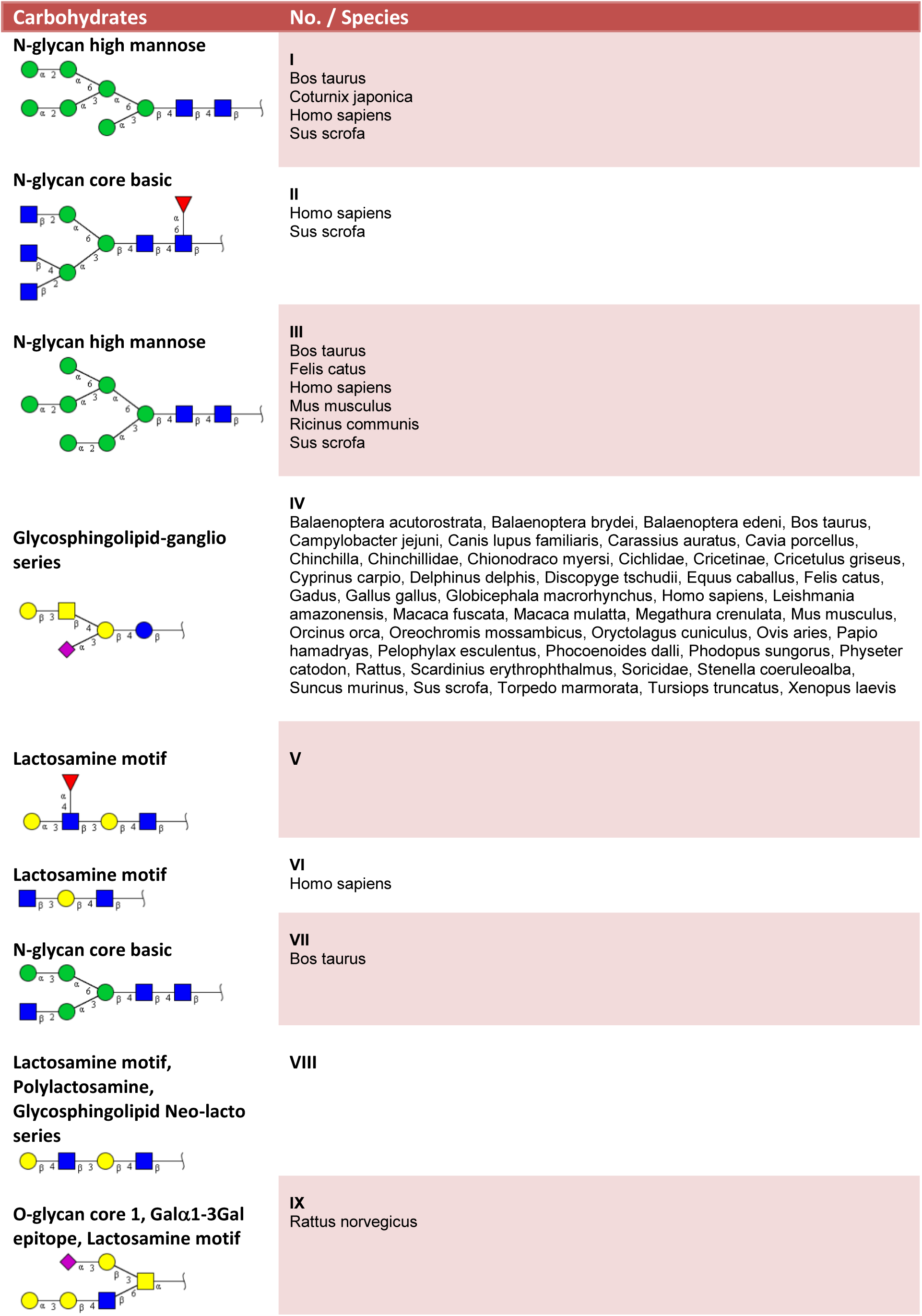
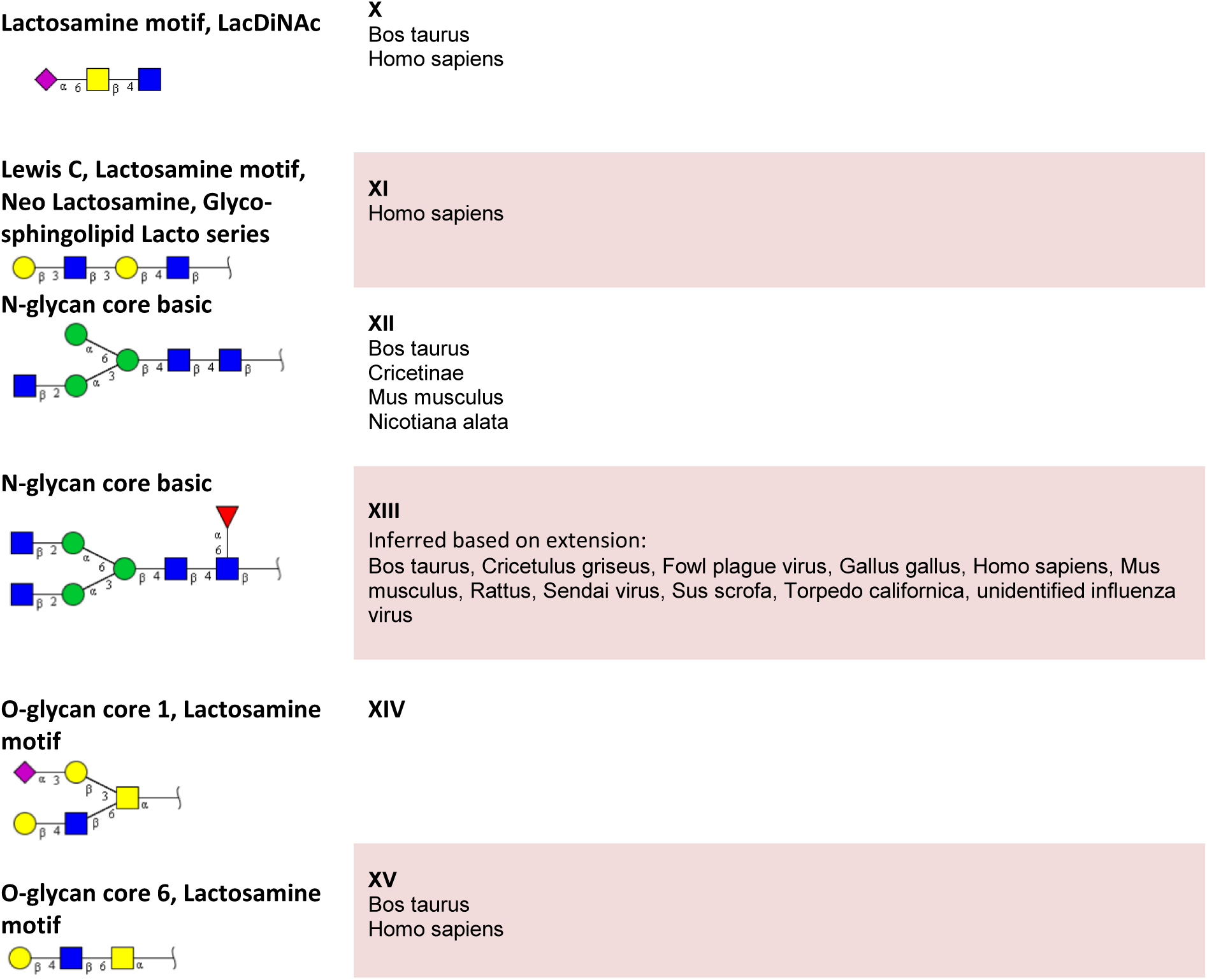
Information on glycans and expression. Species shown *(Species)* were collected from Glycosciences.de websources using MCS. Glycan top-hits were scored in virtual binding to VIP36 conformer 2DUR-A. The top 15 are presented similarly in Figs. S1 and S3. Sequential labeling is in roman numbers.

**Table S2.**
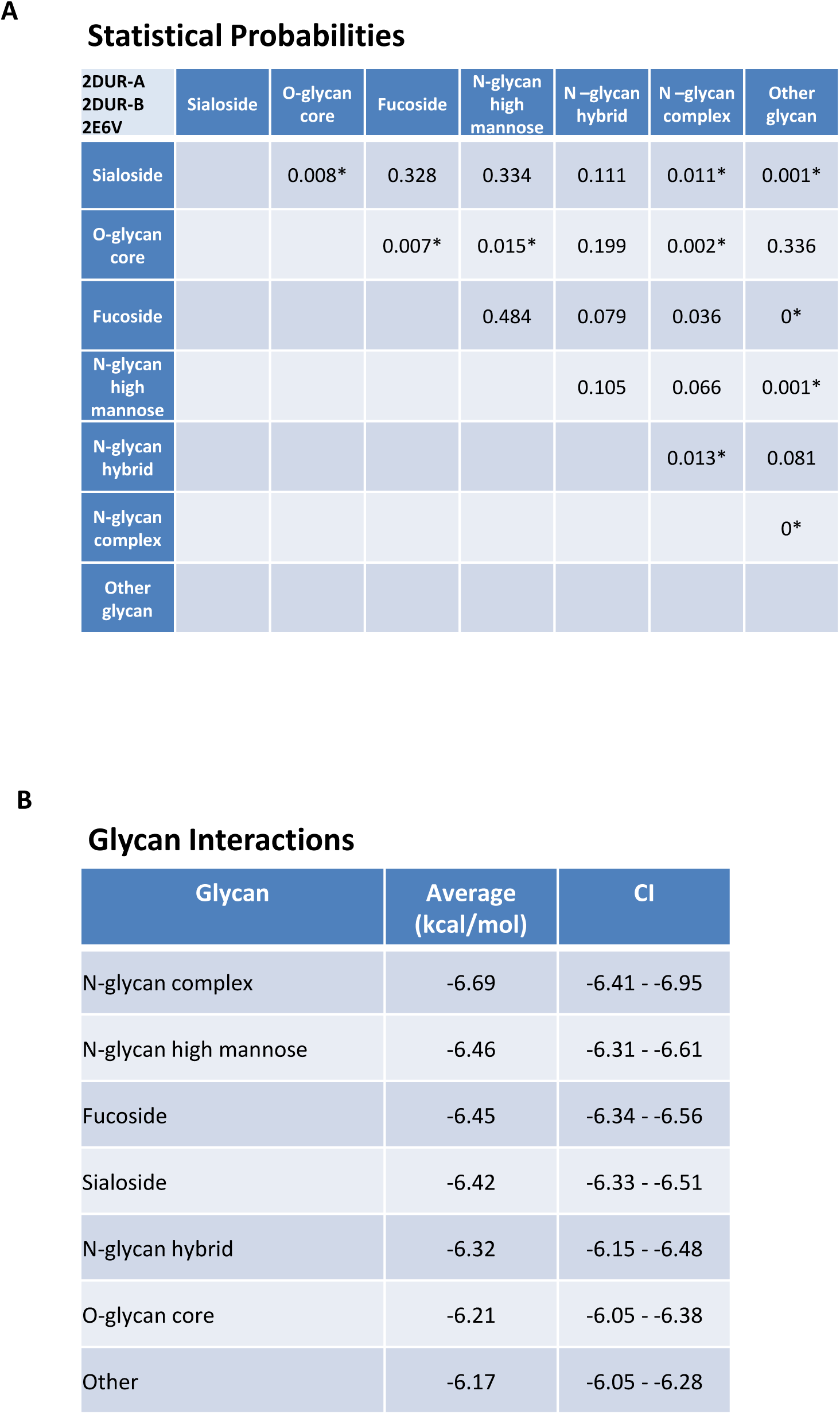
Conformers of 2E6V, 2DUR-A and 2DUR-B were compared for carbohydrate binding in virtual screening with the Carb165 library. **A** T-test with ligands of 2E6V, 2DUR-A and 2DUR-B (n=159-3). The glycans grouped for comparisons were subjected to one-sided T-test. Statistical significances of group differences are plotted. Post-hoc tests show that the values are normally distributed at p>0.08 levels (Kolmogorov-Smirnov and Lilliefors). A Levene statistics of 3.98 indicates the possibility of inflation of error (N=477). For comparison, p values are marked with * if at p<0.05 level with tests with unequal variances; these only showed differences with Fucosides. **B** The average of binding energies to the 2E6V, 2DUR-A and 2DUR-B combined is listed. The confidence interval (CI) is indicated.

**Table S3.**
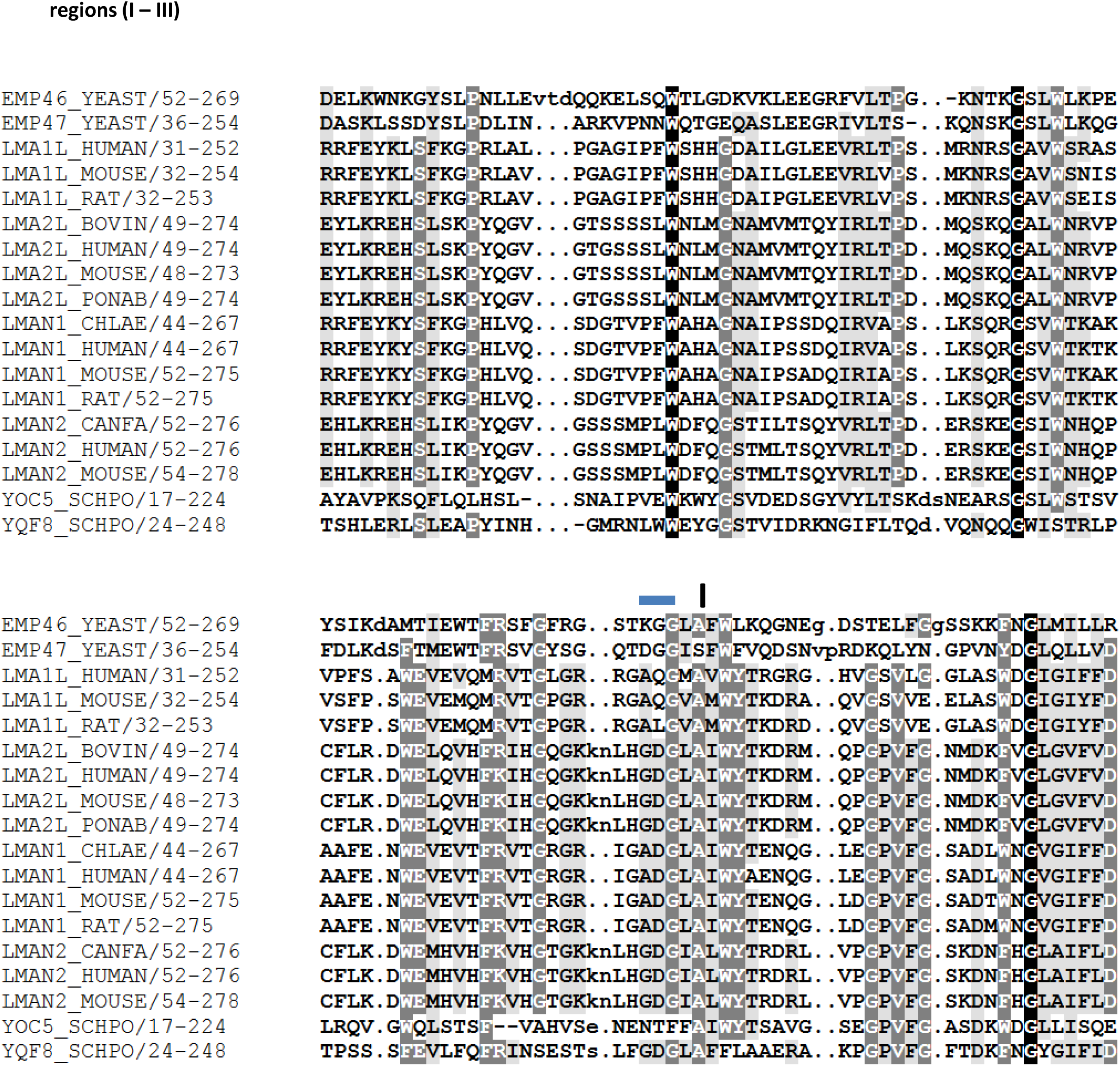
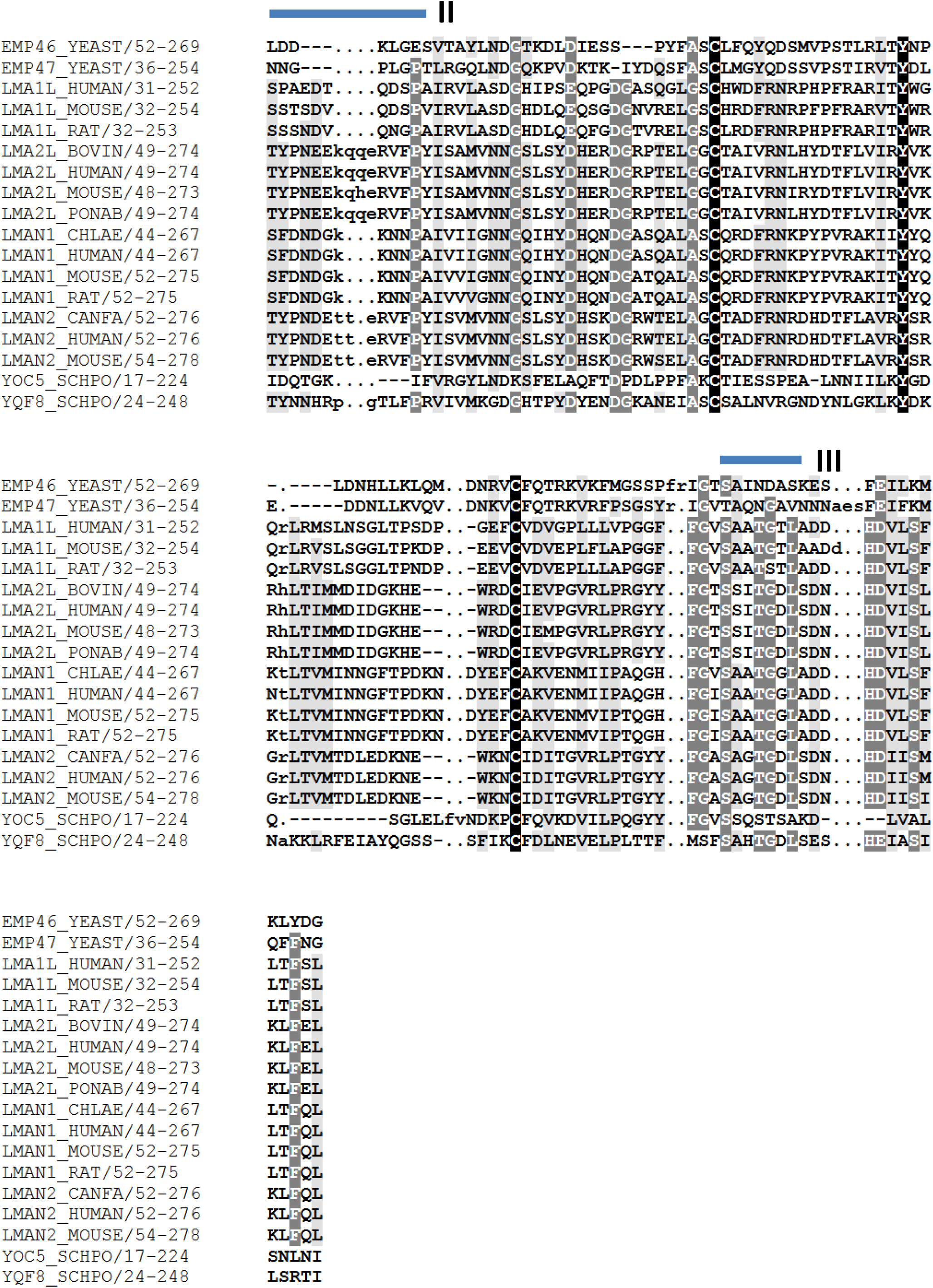
Clustal (Higgins et al 1996) was used to align the Prosite entries (Sigrist et al 2002) that were retrieved with the *Canis familiaris* VIP36 protein. Regions previously described in comparison to leguminous plant lectin Lec_Baupu are labeled I-III.

